# Identification of α-azacyclic acetamide-based inhibitors of *P. falciparum* Na^+^ pump (*Pf*ATP4) with fast-killing asexual blood-stage antimalarial activity by phenotypic screening

**DOI:** 10.1101/2025.05.20.655166

**Authors:** Arturo Casas, Leah S. Imlay, Vandana Thathy, Kate J. Fairhurst, Adele M. Lehane, Aloysus K. Lawong, Ionna Deni, Josefine Striepen, Seungheon Lee, Ashwani Kumar, Chao Xing, Hanspeter Niederstrasser, Bruce A. Posner, Benoît Laleu, Susan A. Charman, David A. Fidock, Joseph M. Ready, Margaret A. Phillips

## Abstract

Malaria treatments are compromised by drug resistance, creating an urgent need to discover new drugs. We used a phenotypic high-throughput screening (HTS) platform to identify new antimalarials, uncovering three related pyrrole-, indole-, and indoline-based series with a shared α-azacyclic acetamide core. These compounds showed fast-killing activity on asexual blood-stage *Plasmodium falciparum* parasites, were not cytotoxic, and disrupted parasite intracellular pH and Na^+^ regulation similarly to cipargamin (KAE609), a clinically advanced inhibitor of the *P. falciparum* Na^+^ pump (*Pf*ATP4). *Pf*ATP4 is localized to the parasite plasma membrane and is essential for maintaining a low cytosolic Na^+^ concentration. Resistance selections on *P. falciparum* parasites with two α-azacyclic acetamide analogs identified mutations in *Pf*ATP4, and cross-resistance was observed across the α-azacyclic acetamides and KAE609, confirming *Pf*ATP4 as the target. *Pf*ATP4 is a well-established antimalarial target, and identification of additional *Pf*ATP4 inhibitors provides alternative avenues to disrupt its function.

## INTRODUCTION

Malaria remains one of the deadliest infectious diseases globally. The World Health Organization (WHO) estimates there were 263 million malaria infections and ∼0.6 million deaths in 2024, with most deaths occurring among children in Africa.^1^ Since 2000, the use of artemisinin (ART)-based combination therapies (ACTs), insecticide-treated bed nets, and other control measures has decreased malaria cases and deaths. However, the decline in malaria deaths has recently stalled. The progress achieved through widespread ACT use is now threatened by emerging drug resistance to both ART and some combination partners. Mutant *k13* alleles that mediate ART partial resistance were first identified in Southeast Asia but have also been found more recently in Africa, the world’s most vulnerable region.^2–5^ Other therapies, such as the RTS,S and R21/Matrix-M vaccines, have been recommended by the WHO for the prevention of *P. falciparum* malaria in young children and can mitigate the severity of infection in some cases, but drug therapy remains essential to malaria treatment and control programs.^1, 6^ As a result, the discovery of drugs and new targets to combat emerging drug resistance and to sustain progress in reducing malaria mortality is imperative.

Malaria is caused by several species of *Plasmodium* parasites, which are transmitted by infected mosquitoes.^6–9^ *Plasmodium falciparum* is responsible for most of the severe cases and deaths, while *P. vivax* has a dormant liver stage that complicates treatment. The malaria parasite’s complex life cycle presents three opportunities for pharmacological intervention including the liver stage that is the site of the initial infection, the intraerythrocytic asexual blood stage (ABS) that cause symptomatic disease, and the intraerythrocytic sexual stage required for transmission. Drugs that treat symptomatic malaria require intraerythrocytic ABS activity, while blocking the liver stage would have prophylactic activity, and gametocyte targeting compounds block transmission.^10^ Fast-acting compounds such as ART derivatives (notably artesunate) provide rapid relief of symptoms for severe malaria cases and may provide reduced resistance risk in the clinic.^9, 11–12^

Most antimalarials in development have emerged from phenotypic high-throughput screens against the ABS parasites.^13–14^ Recently, screens have also yielded compounds with liver-stage and transmission-blocking activity.^15–16^ Target-based approaches have also been successful in producing clinical candidates. For instance, a dihydroorotate dehydrogenase (DHODH) enzyme-based high-throughput screen (HTS) led to the discovery of DSM265, which reached Phase IIa clinical development.^17^ An advantage of phenotypic screening is that it selects for compounds with inherent cell permeability, although identifying their molecular targets can be challenging. In malaria drug discovery programs, resistance screening followed by whole-genome sequencing (WGS) has been a robust mechanism to identify the targets of many compounds.^18^ This approach successfully identified the targets of several antimalarial candidates currently in development, including the Phase II clinical candidate cipargamin (KAE609).^19–23^ KAE609 targets a plasma membrane ATP-dependent transporter (*Pf*ATP4) that is essential to maintain a low cytosolic Na^+^ concentration and that is believed to import H^+^ while extruding Na^+^.^24–25^ A second clinical candidate, SJ733, which completed a Phase IIa study in Peru, has also been shown to act on PfATP4.^26–27^

Herein, we describe four structurally related compounds that have a common α-azacyclic acetamide pharmacophore: pyrroloacetamides **1** (SW412) and **2** (SW491), and indoloacetamides **3** (SW968), and **4** (SW080) (Figure 1A). All four compounds have fast-killing activity against ABS *P. falciparum*. These compounds were discovered in a previously reported phenotypic HTS for small molecule compounds with *P. falciparum* ABS antimalarial activity. This screen also identified a tetrazole series targeting heme polymerization^28^, a tyrosine-amide series that linked to *Pf*CARL-mediated resistance^29^, and a piperidine carboxamide series targeting the proteasome β5 subunit.^30^ Treatment of *P. falciparum* parasites with **1**-**3** disrupted parasite pH and Na^+^ regulation, leading to an increase in intracellular Na^+^ and pH, similar to KAE609. Drug selections for **2**- and **3**-resistant parasites followed by WGS showed that resistance was linked to mutations in *Pf*ATP4, further confirming the target of these compounds. Consistent with these findings, cross-resistance was observed between KAE609 and the α-azacyclic acetamides from our screen, indicating that all five compounds share *Pf*ATP4 as a common target. The similar drug-like properties of the α-azacyclic acetamides make them appealing targets for medicinal chemistry if additional *Pf*ATP4 compounds are needed in the antimalarial portfolio.

**Figure 1.**
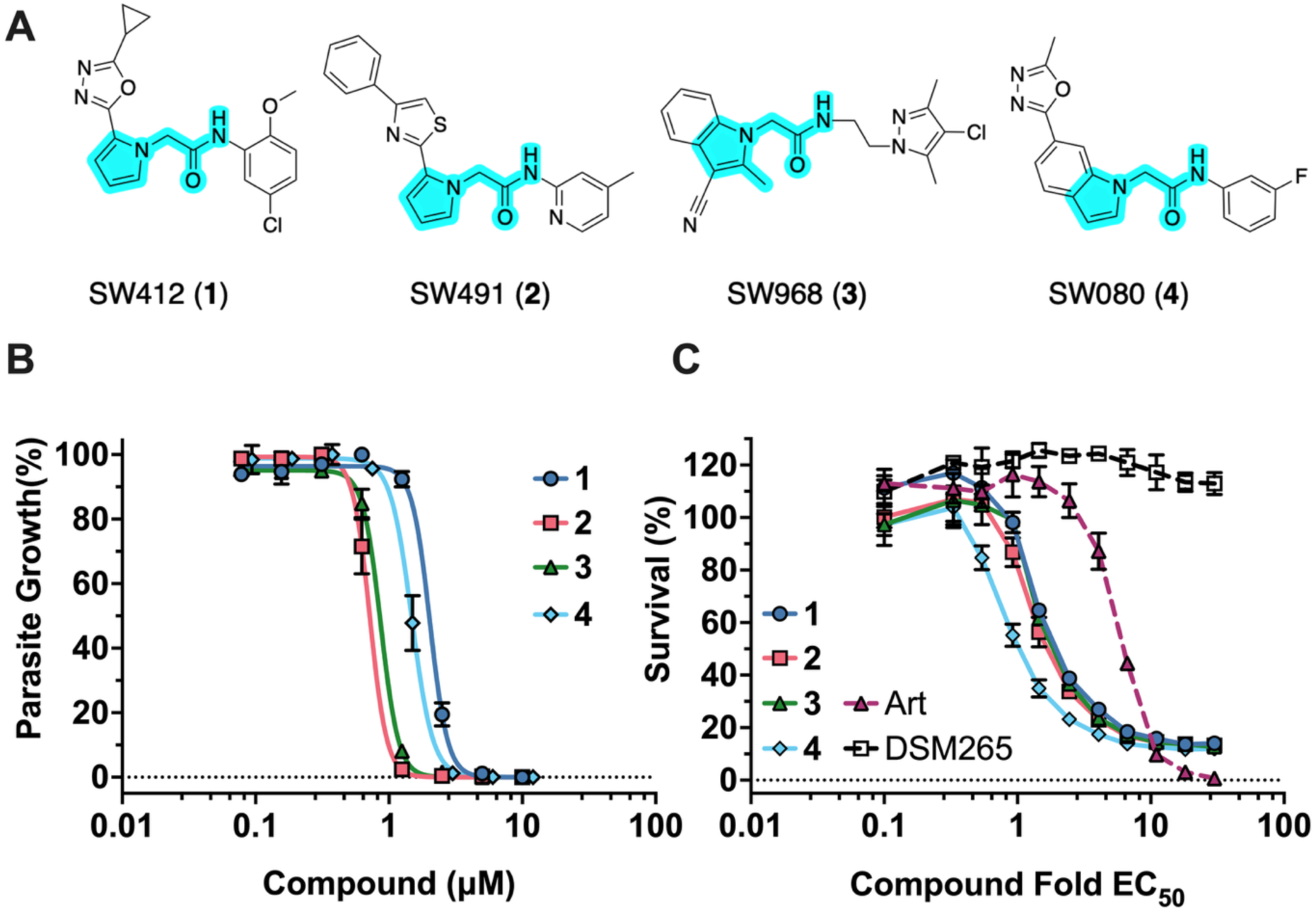
Potency and kill rate of α-azacyclic acetamides versus *P. falciparum* asexual blood stage (ABS) parasites. (A) Chemical structures of **1**-**4**. The common α-azacyclic acetamide structural motif is highlighted in blue. (B) *P. falciparum* (strain Dd2) growth inhibition (ABS parasites) by the α-azacyclic acetamides (**1**-**4**). Representative data from one EC_50_ experimental trial (3 technical replicates) showing the mean ± std dev. See Table 1 for mean EC_50_ data for multiple independent studies per compound. (C) Kill rate was assessed using the BRRoK (bioluminescent relative rate of kill) assay. *P. falciparum* NF54^luc^ parasites were incubated for 6 h with **1**-**4** and control compounds artemisinin (ART; Fast Kill) and DSM265 (Slow Kill), over a concentration range from 30-0.10x the 72 h *Pf*3D7 EC_50_ (average EC_50_ values used for these studies were **1** (2.1 μM), **2** (0.70 μM), **3** (0.92 μM), **4** (1.9 μM), ART (0.013 μM) and DSM265 (0.0080 μM)). A representative data set is shown, and data are the mean ± std dev for 3 technical replicates. Data for a second independent experiment are shown in Figure S1.

**Table 1.** Summary of potency and *in vitro* ADME data on the α-azacyclic acetamides.

|  | **1** | **2** | **3** | **4** |
| --- | --- | --- | --- | --- |
| Activity of all compounds on *Pf* and human cells (μM) | | | | |
| *Pf*3D7 EC<sub>50</sub> | 2.5 ± 0.26 (5) | 0.83 ± 0.071 (5) | 1.1 ± 0.071 (5) | 2.<u>0</u> ± 0.38 (3) |
| *Pf*3D7 pH EC<sub>50</sub> | 4.5 ± 2.<u>0</u> (2) | 1.6 ± 0.95 (2) | 1.5 ± 0.047 (2) | N/A |
| *Pf*Dd2 EC<sub>50</sub> | 2.4 ± 0.13 (15) | 0.83 ± 0.048 (16) | 1.1 ± 0.072 (16) | 1.6 ± 0.1<u>0</u> (8) |
| *Pf*NF54<sup>luc</sup> EC<sub>50</sub> | 3.0 ± 0.93 (2) | 0.72 ± 0.019 (2) | 0.94 ± 0.0027 (2) | 1.2 ± 0.0055 (2) |
| *Pf*D10 attB EC<sub>50</sub> | 2.9 (1) | 1.2 (1) | 1.3 (1) | 1.7 (1) |
| *Pf*D10 attB + yDHODH EC<sub>50</sub> | 3.2 (1) | 0.99 (1) | 0.93 (1) | 1.9 (1) |
| Human HepG2 CC<sub>50</sub> | > 50 (3) | > 50 (3) | > 50 (2) | > 50 (2) |
| Chemical properties |  |  |  |  |
| Molecular weight | 372 | 374 | 369 | 350 |
| Predicted pKa | Basic: None<br>Acidic: 11.7 | Basic: 4.8<br>Acidic: 11.8 | Basic 3.1<br>Acidic None | Basic None<br>Acidic None |
| cLogD (pH7.4) | 2.5 | 4.7 | 2.4 | 2.5 |
| Solubility and metabolic stability |  |  |  |  |
| Solubility (pH 2.0) μM | 25 | 99 | 25 | 25 |
| Solubility (pH 6.5) μM | 12.5 | 3.2 | 25 | 25 |
| Human CL<sub>int</sub>, *in vitro* (μL/min/mg protein) | 279 | >866 | >866 | 126 |

## RESULTS

### Identification of hit compounds with a common α-azacyclic acetamide pharmacophore

We previously described a screen of a compound library consisting of 100K small molecules against ABS *P. falciparum* 3D7 parasites (*Pf*3D7).^28–30^ Several unreported antimalarial compounds from the screen also showed sub-micromolar activity, while not showing cytotoxicity. From this set, four additional compounds were selected for mechanism of action (MOA) studies based on a common α-azacyclic acetamide pharmacophore, including **1**-**4** (Figure 1A). These compounds had low-micromolar activity against both drug-sensitive *Pf*3D7 and drug-resistant (chloroquine, pyrimethamine, and low-level mefloquine) *P. falciparum* Dd2 (*Pf*Dd2) parasites (Figure 1B and Table 1), while showing no cytotoxicity versus a human hepatic cell line (HepG2 CC_50_ (cytotoxic concentration to reach 50%) > 50 μM) (Table 1). Similar activity was noted for all compounds against a *P. falciparum* strain expressing *Saccharomyces cerevisiae* yeast DHODH (*Pf*D10 attB + yDHODH) (Table 1), ruling out mitochondrial DHODH and/or bc1 complex as potential targets.^31–32^ All four compounds have drug-like properties according to Lipinski’s rules.^33^ In early ADME (absorption, distribution, metabolism, and excretion) studies, all except **2** showed reasonable kinetic solubility in phosphate buffer at pH 6.5 (Table 1). Metabolic stability in human liver microsomes was poor, indicative of early hit compounds that require optimization to obtain an early lead compound.

To establish preliminary structure-activity relationships (SAR), a small set of additional analogs of **2** and **3** were purchased from commercial sources and evaluated for activity against *P. falciparum* 3D7 and Dd2 parasites (Table S1). Analogs of **2** probed the impact of moving the meta-methyl on the pyridine ring to the para position (SW463; **5**), of moving the pyridine nitrogen ortho to the methyl (SW316; **6**) or of adding a *para*-chloro substituent to the benzyl ring (SW317; **7**). These modifications were detrimental, leading to 10-20-fold reduction of activity. Combining the ortho-methyl with the para-chloro benzyl (SW318; **8**) led to a complete loss of activity. Replacement of the pyridine nitrogen with carbon (SW319; **9**) led to a 2-fold reduction of activity, though activity was restored in the context of a chloro substitution for the meta-methyl (SW320; **10**). Replacement of the substituted pyrrole with indoline led to a complete loss of activity (SW966; **11**). Only a small set of analogs to **3** were available, and these analogs assessed removal of the indole methyl (SW181; **12**), removal of the acetyl group (SW315; **13**), or both (SW314; **14**), all of which led to loss of activity (Table S1). The significant changes in activity observed with relatively modest modifications to the chemical structures of these analogs strongly suggest that the alterations impact binding interactions with a specific protein target, rather than common mechanisms such as hemozoin binding or reactive oxygen species generation.

### The α-azacyclic acetamides show fast-killing activity

Compounds with fast kill rates are of particular interest because they can provide rapid relief of malaria symptoms.^11–12^ α-azacyclic acetamides **1**-**4** were tested to determine their kill rate using a previously described bioluminescent relative rate of kill (BRRoK) assay that evaluates the ability of a compound to block growth over a very short assay window of 6 h.^34^ Briefly, following a 6 h exposure to each compound over a concentration range from 30 - 0.10 times the EC_50_, transgenic NF54^luc^ parasites were evaluated for growth based on a luciferase assay system as the readout. The α-azacyclic acetamides were benchmarked alongside reported fast kill (artemisinin;ART) and slow kill (DHODH inhibitor, DSM265) compounds^17, 35^ (Figures 1C and S1). The α-azacyclic acetamides **1**-**4** showed similar profiles to ART in this assay supporting the conclusion that they have a fast kill mechanism. Interestingly, the α-azacyclic acetamides induced killing at a lower concentration relative to EC_50_ than for ART. The slow killing control DSM265 did not reduce parasitemia in the 6 h treatment window of the experiment, as expected.

### The α-azacyclic acetamides cause an increase in cytosolic pH consistent with PfATP4 inhibition

Prior studies have established that the MOA of antimalarial compounds is correlated with compound kill rate.^36 34^ Many compounds with fast kill MOAs have been found to inhibit the plasma membrane ATP-dependent transporter *Pf*ATP4, including a significant percentage of compounds in the Medicines for Malaria Venture’s (MMV) Malaria Box and Pathogen Box.^37–38^ Thus, as the next step to identify the MOA of α-azacyclic acetamides we conducted cytosolic pH assays to determine if they could be *Pf*ATP4 inhibitors. *Pf*ATP4 functions as a Na^+^ efflux pump that also drives the counter-flow of protons into the cell (Figure 2A).^19, 24–25, 39–40^ Compounds targeting *Pf*ATP4 cause an increase in parasite cytosolic Na^+^ concentration ([Na^+^]_cyt_), leading to swelling of the parasite and the infected red blood cell (RBC), as well as an increase in cytosolic pH (pH_cyt_).

**Figure 2.**
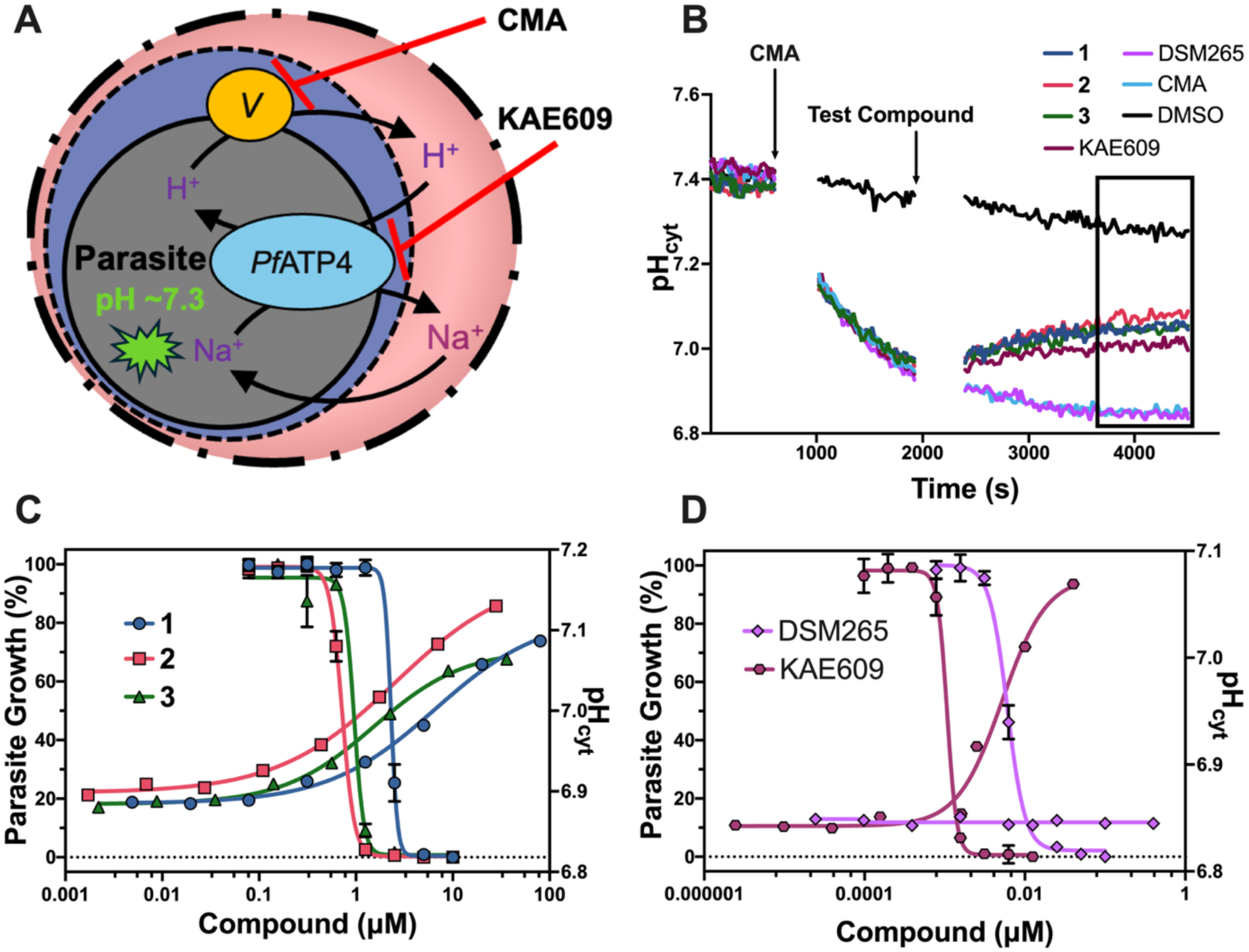
Impact of the α-azacyclic acetamides **1**-**3** on intracellular pH of *P. falciparum* 3D7 parasites. (A) Schematic diagram illustrating the interplay between ion transport mechanisms in the malaria parasite: The Na^+^-ATPase *Pf*ATP4 exports Na^+^ ions while simultaneously importing H^+^ ions and the V-type H^+^-ATPase (V) provides an opposition to acidification. Concanamycin A (CMA), a V-type inhibitor causes a decrease in intracellular pH, while administration of *Pf*ATP4 inhibitors (e.g. KAE609) leads to alkalinization. This diagram features the host RBC (salmon compartment), the parasitophorous vacuole (indigo compartment), and the parasite (charcoal compartment). (B) Parasite pH_cyt_ over time showing the effects of α-azacyclic acetamides **1**-**3** and control compounds. *Pf*3D7 trophozoites isolated by saponin lysis were loaded with the pH-sensitive fluorescent dye BCECF. CMA (50 nM) was added at 10-min to acidify the cytosol via inhibition of the V-type H^+^-ATPase. A DMSO-only (1% v/v) control without CMA was also included in the study (black trace). Compounds (including a DMSO-only control – turquoise trace) were added at 30-min, and the impact on cytosolic pH was monitored. Representative data from one experimental trial are shown. Compounds were tested at 10× EC_50_ based on the SYBR green 72 h assay: **1** (20 μM), **2** (7 μM), **3** (9 μM), DSM265 (0.1 μM; negative control) and KAE609 (0.01 μM; positive control) with experimental concentrations shown in parenthesis. A second independent replicate of this experiment is shown in Figure S2B. (C) The average pH_cyt_ value obtained from the final 10-minute period of the assay-corresponding to a time window when the maximal CMA-induced acidification is observed in the controls DSM265 and CMA (see boxed 10-minute period in part B) is plotted on the right-hand axis. This time matched window was used to compare responses to α-azacyclic acetamides **1**-**3** and KAE609, over 8 concentrations ranging from 40 - 0.002x EC_50_. pH data represent a single study at n=1 per concentration. A second independent replicate of the pH study is provided in Figure S2C. The corresponding *P. falciparum* (strain 3D7) growth inhibition data are plotted (left-hand axis) over a range of compound concentrations as measured by the SYBR green 72 h assay. Data represent the mean ± std dev for one representative EC_50_ experimental trial (3 technical replicates). Table 1 shows the average data from at least three independent studies per compound. pH vs time curves for all drug concentrations that support this plot are shown in Figure S3. (D) Similar graphical representation and EC_50_ overlay as previously discussed in (C) for controls DSM265 and KAE609. A second independent replicate of the pH arm of this study is provided in Figure S2D.

A cell-based assay for ATP4 inhibition has been reported that monitors intracellular pH (pH_cyt_).^24, 40–41^ In this assay, the pH responses of isolated malaria parasites following exposure to compounds of interest are measured in the presence of the V-type H^+^-ATPase inhibitor concanamycin A (CMA) (Figure 2A). V-type H^+^-ATPase is a parasite plasma membrane protein that contributes to the maintenance of cytosolic pH in the range of 7.2–7.3, functioning in the opposite direction to ATP4 to pump protons back out of the cell.^42–44^ Inhibition of V-type H^+^ ATPase leads to acidification of the parasite cytosol, while inhibitors of *Pf*ATP4 in contrast cause alkalinization of the parasite cytosol. The effects of PfATP4 inhibitors on pH_cyt_ have been shown to be more readily detected when the V-type H^+^ ATPase is inhibited with CMA.

α-azacyclic acetamides **1**-**3** were tested in this assay on *P. falciparum* 3D7 parasites alongside the known *Pf*ATP4 inhibitor KAE609 as a positive control and the DHODH inhibitor DSM265 as a negative control. An increase in pH_cyt_ was observed in the presence of both KAE609 and the three tested α-azacyclic acetamides, suggesting that they too are *Pf*ATP4 inhibitors (Figures 2B and S2B). As expected, the addition of 1% v/v DMSO or DSM265 did not lead to alkalinization of the parasite cytosol. The DMSO-only control in the absence of CMA showed no pH increase. To interrogate the relationship between intracellular pH changes and the cell-killing activity of α-azacyclic acetamides **1**-**3** we conducted the pH assay over a range of compound concentrations. The intracellular pH was averaged over a ten-minute period to determine a steady-state pH after compound addition (Figures 2B, S2B, S3 and S4 black box) and this value was plotted versus compound concentration for **1**-**3** (Figures 2C and S2C) and for control compounds KAE609 and DSM265 (Figures 2D and S2D). The pH data were overlaid with the compounds impact on parasite growth measured in the standard 72 h SYBR Green assay across the same concentration range. This analysis showed that the 50% (EC_50_) effective concentration to cause alkalinization of the parasite cytosol in the pH assay was similar (within 2-fold) to the growth inhibition EC_50_, providing strong evidence that the ionic dysregulation was linked to cell death (Tables 1 and S2). Similarly, the EC_50_ for KAE609 in the growth assay was within 3.2-fold of the pH EC_50_, suggesting that the compound’s impact on ion homeostasis directly correlates with cell killing. In contrast, for DSM265, there was no impact on pH and therefore, no correlation with cell killing.

### Testing of α-azacyclic acetamides in a pH fingerprint assay provides further evidence for PfATP4 inhibition

α-azacyclic acetamides **2** and **3** were additionally characterized in a pH fingerprint assay, which can detect (and discriminate between) protonophores and inhibitors of a number of plasma membrane transporters including *Pf*ATP4, the lactate/H^+^ transporter *Pf*FNT, the V-type H^+^-ATPase, the hexose transporter *Pf*HT, and the acid-loading Cl^−^ transporter(s).^41^ In this assay, parasites were isolated from their host erythrocytes, loaded with the pH-sensitive dye BCECF, and depleted of ATP by incubation in a glucose-free solution. Parasites were then placed in three separate solutions, and their pH response monitored over time. The α-azacyclic acetamides **2** and **3** gave rise to a ‘pH fingerprint’ consistent with *Pf*ATP4 inhibition (Figure S5). When tested at 5 μM, **2** had a similar impact on the intracellular pH as KAE609 at 50 nM (a concentration that causes maximal inhibition of *Pf*ATP4), whereas **3** gave rise to an intermediate phenotype between those of KAE609 and the solvent control in the condition (+Glucose +CMA) in which *Pf*ATP4 inhibitors can be detected (Figure S5B). These results are consistent with 5 μM **2** having caused maximal/near-maximal inhibition of *Pf*ATP4, and **3** sub-maximal inhibition. Neither **2** nor **3** showed evidence of having any of the other MOAs that could be detected with this assay (Figure S5).

### The α-azacyclic acetamides cause accumulation of intracellular Na+ consistent with inhibition of PfATP4

After the invasion by the malaria parasite (Figure 3A), for 12-18h (ring stage), the permeability of the RBC membrane increases significantly, allowing an influx of Na^+^ and other small solutes.^45^ Na^+^ enters the infected RBC down its concentration gradient via new permeability pathways induced by the parasite. The parasitophorous vacuolar membrane is believed to be freely permeable to small solutes like Na^+^ (trophozoite stage). Therefore, the Na^+^ concentration within the parasitophorous vacuole is expected to be like that in the RBC cytosol. However, despite the high Na^+^ concentration in its surrounding environment, the parasite maintains a low intracellular Na^+^ concentration ([Na^+^]_cyt_) by actively extruding Na^+^ through *Pf*ATP4.^24, 46^ Consistent with this mechanism, *Pf*ATP4 inhibitors cause significant Na^+^ accumulation within the cytosol.^24^ α-azacyclic acetamides **2** and **3** were tested to determine if they also caused an increase in [Na^+^]_cyt_ at two different concentrations, with KAE609 and solvent alone (0.1% v/v DMSO) included as controls (Figure 3B). As expected, KAE609 and the a-azacyclic acetamides gave rise to an increase in [Na^+^]_cyt_. When tested at 5 mM, both **2** and **3** showed a similar increase in cytoplasmic Na^+^ to a supramaximal concentration of KAE609 (50 nM), whereas at 1 mM, **2** and **3** had a reduced response, consistent with their lower potency against the parasite in proliferation assays. As expected, the presence of 0.1% v/v DMSO did not lead to an increase in [Na^+^]_cyt_.

**Figure 3.**
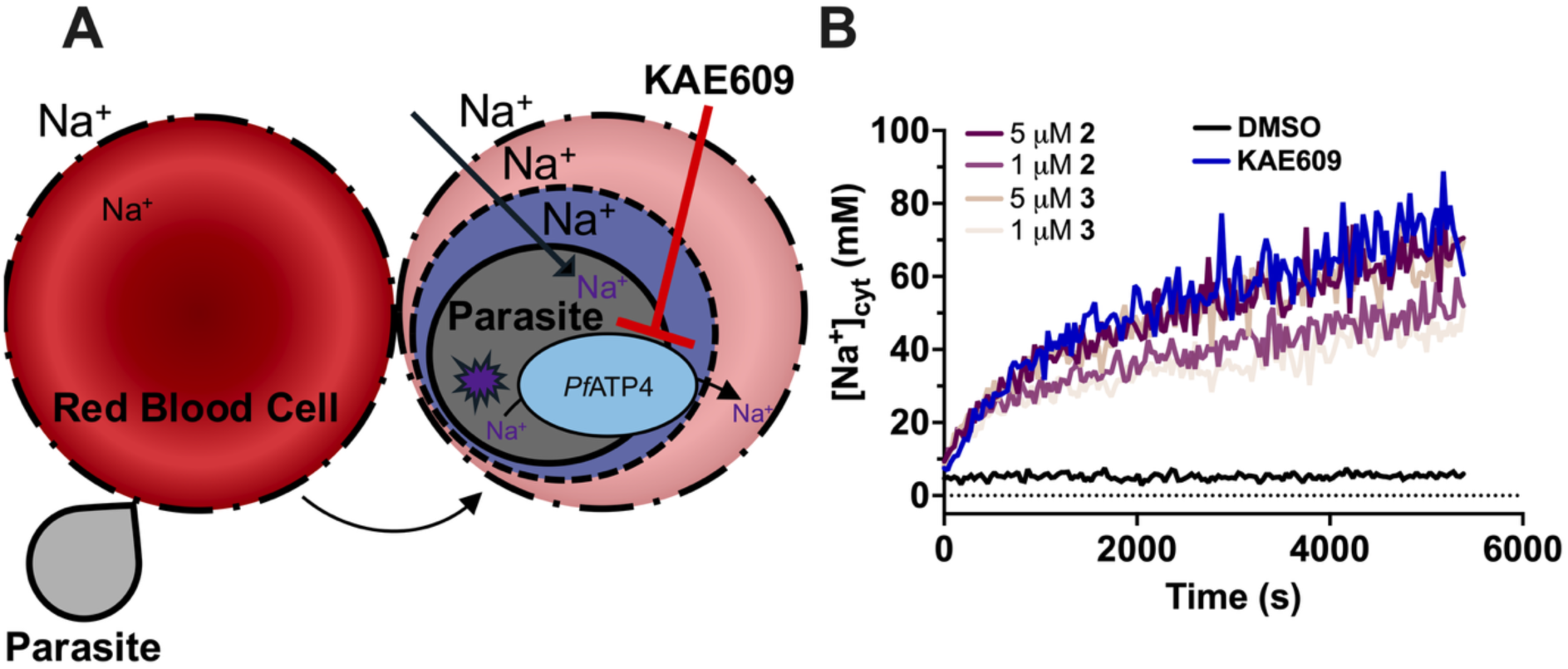
Impact of the α-azacyclic acetamides on intracellular Na^+^ concentration (A) Schematic diagram illustrating the inward Na^+^ electrochemical gradient across the plasma membrane of an intraerythrocytic trophozoite, maintained by the Na^+^-ATPase, *Pf*ATP4, which exports Na^+^ ions from the parasite cytosol. Administration of *Pf*ATP4 inhibitors (e.g. KAE609) leads to accumulation of Na^+^ ions in the parasite cytosol. The diagram features the host RBC pre-invasion (red) alongside a post-invasion RBC (salmon compartment) surrounded by its membrane (broken line), the parasitophorous vacuole (indigo compartment) enclosed by its membrane (dashed line), and the parasite pre-invasion (grey) alongside a post-invasion parasite (charcoal compartment) surrounded by its plasma membrane (solid line). (B) The effects of **2** (1 μM and 5 μM), **3** (1 μM and 5 μM), KAE609 (50 nM; positive control for PfATP4 inhibition), and DMSO (0.1% v/v; solvent control) on [Na^+^]_cyt_. The experiment was performed with isolated *Pf*3D7 trophozoites loaded with the Na^+^-sensitive fluorescent dye SBFI. The data are from a single experiment, representative of two similar experiments in which **2** and **3** were tested.

### Selection of 2- and 3-resistant PfDd2 parasites

To confirm the MOA for the α-azacyclic acetamides against *P. falciparum* and to obtain potential information about whether or not they bind *Pf*ATP4 at an overlapping site with that of KAE609, we undertook resistance selections using *Pf*Dd2 parasites using the published cycling drug selection method.^11, 47^ α-azacyclic acetamides **2** and **3** were chosen as representative examples of the pyrrole- and indole-based series, respectively. Parasites (∼10^9^/flask; 4 flasks per compound) were treated with compound (10×EC_50_) for 48 h, which was sufficient to clear parasites to below the level of detection, and then cultures were allowed to recover in the absence of drug until parasites recrudesced (Table S3). Additional cycles were performed at 20×EC_50_, until resistant parasites were detected based on a fold change in EC_50_ compared to the parental clonal Dd2 line (Table S3). Resistant parasites were detected in the **2**-selections in all 4 flasks after 4 pulses and for **3**-selections in 3 flasks after 5 pulses. Clonal lines were generated from each flask, and one clone per flask was selected for WGS and EC_50_ determination (Tables S4 and S5). WGS analysis identified two single point mutations in *Pf*ATP4 that were not present in the original clonal Dd2 parasite line (Tables 2 and S4). P412L was found in three of four flasks containing **2**-selected resistance clones, and F917L was identified in all three flasks containing **3**-selected clones and in the fourth flask containing **2**-selected clones (Table S4). *Pf*ATP4 mutations identified by the WGS analysis in parasite lines were verified by PCR of the *pfatp4* gene, followed by Sanger sequencing (Figures S6, S7 and Table S6). An additional protein coding gene (ribosomal protein S8e) was mutated in the **2**-selections. However, this mutation was identified in only one of four sequenced clones and is therefore unlikely to be related to resistance to **2** (Figure S8). All three resistant clones derived from the **3**-selections harbored an identical mutation in a gene (*pfcpu)* encoding a conserved protein of unknown function, which was absent in the parental line (Figure S9). The *pfmdr1* locus showed a mixed sequence at position 86 (A or T) in both the parental and resistant cell lines from **3**-selections (Figure S10) consistent with parasites harboring multiple copies of the gene with sequence differences at this position.

One clonal line harboring each of the two identified *Pf*ATP4 mutations (P412L and F917L) was further profiled for susceptibility to **2** and **3**, and tested for cross-resistance to **1**, **4**, and KAE609 (Figure 4A and 4B). EC_50_ increases ranged from 3.4 to 35-fold (Tables 2, S5 and Figure 4C-G). The P412L mutation was associated with the largest EC_50_ increases against the compound used for its selection (**2**), whereas the F917L mutation was associated with relatively high levels of resistance towards all 5 compounds.

**Figure 4.**
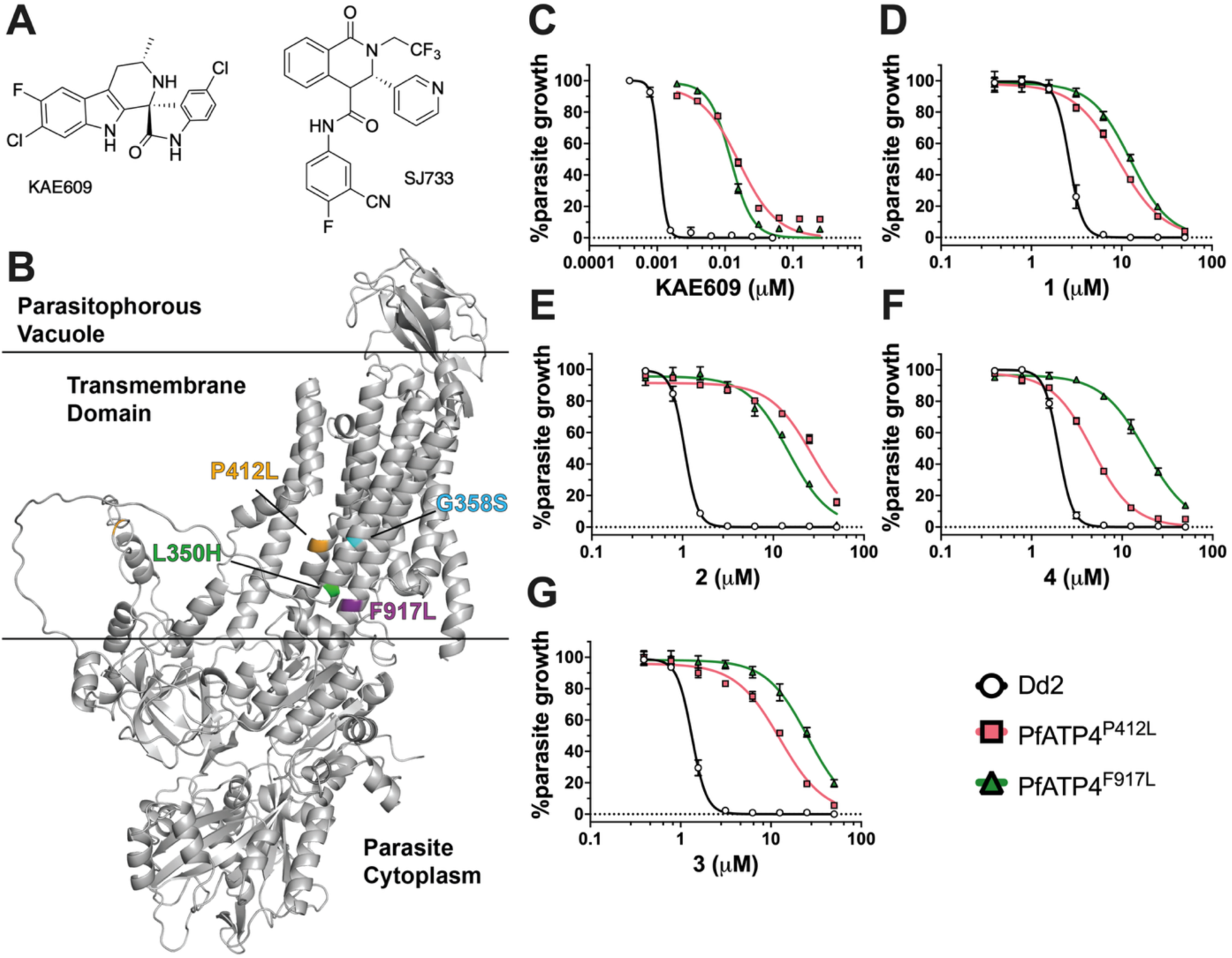
Evaluation of cross-resistance between the α-azacyclic acetamides and KAE609 in parasites with wild-type *Pf*ATP4 mutations associated with resistance to **2** and **3** (A) Chemical structures of clinical candidates KAE609 and SJ733. (B) AlphaFold-derived model of the *Pf*ATP4 structure. Mutations analyzed in this study are found in the transmembrane domain and are labeled; their positions are depicted by the coloring of the residue to match the label color. The structure was displayed using PyMol 3.1.3. (C-G) *P. falciparum* dose-response data for C) KAE609, D-G) α-azacyclic acetamides (**1**-**4**) parasite growth assays performed with *Pf*ATP4 P412L and F917L mutant Dd2 cell lines. Representative data from one EC_50_ determination (3 technical replicates) are shown; data points represent mean ± std dev and units are μM. Mean EC_50_ data for multiple independent replicates are provided in Table 2.

**Table 2.** Comparison of EC_50_ values for WT Dd2 and *Pf*ATP4 mutant lines selected for resistance to α-azacyclic acetamides 2 and 3.

| Parasite line | Selection agent | Key mutations | **1**<br>(μM) | **2**<br>(μM) | **3**<br>(μM) | **4**<br>(μM) | KAE609<br>(nM) |
| --- | --- | --- | --- | --- | --- | --- | --- |
| Dd2 | N/A | N/A | 2.4<sup>a</sup> | 0.83<sup>a</sup> | 1.1<sup>a</sup> | 1.6<sup>a</sup> | 1.2 ± 0.087 (9) |
| **2**-selected<br>Flask #3<br>Clone F5 | **2** | *Pf*ATP4<sup>P412L</sup> | 8.2 ± 1.2<br>(3)[3.4] | 29 ± 6.1<br>(3)[35] | 11 ± 1.8<br>(3)[10] | 3.8 ± 0.32<br>(5)[2.4] | 13 ± 1.1<br>(5)[11] |
| **3**-selected<br>Flask #3<br>Clone D8 | **3** | *Pf*ATP4<sup>F917L</sup> | 11 ± 0.54<br>(3)[4.6] | 14 ± 0.79<br>(3)[17] | 27 ± 0.71<br>(3)[25] | 18 ± 0.21<br>(3)[11] | 13 ± 0.53<br>(5)[11] |

<sup>a</sup> Data were reproduced from Table 1. All other data represent the mean EC<sub>50</sub> ± SEM with the number of independent replicates shown in parentheses. Triplicate technical replicates were performed for each individual study. Fold EC<sub>50</sub> changes compared to WT Dd2 are displayed in brackets. Ordinary one-way ANOVA analysis with Dunnett correction was performed in Graphpad Prism 10 for each compound across strains; P-values were <0.0001 for comparison to the WT Dd2 data in all cases. The highest concentrations in the dose-response titrations were 50 μM for compounds **1-4** and 50 - 250 nM for KAE609.

### Cross-resistance with known PfATP4 inhibitors

To further profile common resistance alleles versus previously described *Pf*ATP4 inhibitors, the α-azacyclic acetamides were tested against three additional Dd2 *Pf*ATP4 mutant clones that had been selected for resistance to the dihydroisoquinolone SJ733^27^ or its backup analog MMV1793609 (**15**)^9, 48^ (Figure 4A, Tables 3, S7 and S8). These lines harbored two mutations (G358S and L350H) we did not identify in resistance selections for **2** and **3**, while the third overlapped (P412L). These studies were conducted in the Fidock lab, where the EC_50_ values for **2** and **3** in parental Dd2 parasites were approximately 2.5-fold lower (Table 3) than those observed in the Phillips Lab (Table 2). The *Pf*ATP4^G358S^ mutation originally identified in selections for SJ733 resistance^27^ was also identified in patients in the KAE609 clinical trial and was conferred high-level resistance to both SJ733 and KAE609^21^. Our study similarly found that the G358S mutation large EC_50_ shifts for both SJ733 (240-fold EC_50_ increase) (Table S8) and to KAE609 (approximately 600- to 700-fold EC_50_ increase) (Tables 3 and S8). In contrast, only a 1.9-fold increase was observed against **3**, while a slight decrease in EC_50_ was observed against **2** (Table 3), suggesting that the binding site of the α-azacyclic acetamides may not fully overlap with KAE609 and SJ733. The L350H mutation led to only modest 4- to 6-fold EC_50_ increases for all three compounds (**2**, **3** and KAE609). The Fidock lab-derived *Pf*ATP4^P412L^ line showed EC_50_ increases for all three compounds (Table 3) similar to that observed for the Phillips lab-derived P412L mutant clone (Tables 2 and 3).The *Pf*ATP4 mutations identified in our current study (F917L and P412L) and the tested SJ733/**15**-resistant mutations (L350H and G358S) (Table 3) map to the alpha-fold predicted transmembrane region of *Pf*ATP4 (Figure 4B).

**Table 3.** Susceptibility of lines with SJ733- or 15-selected *Pf*ATP4 mutations to α-azacyclic acetamides 2 and 3 and KAE609.

| Parasite line | Selection Agent | *Pf*ATP4 genotype | **2**<br>EC<sub>50</sub> (μM) | **3**<br>EC<sub>50</sub> (μM) | KAE609<br>EC<sub>50</sub> (nM) |
| --- | --- | --- | --- | --- | --- |
| <sup>a</sup>Dd2-B2 | n/a | *Pf*ATP4<sup>WT</sup> | 0.38 ± 0.031 | 0.54 ± 0.054 | 1.3 ± 0.10 |
| Dd2<sup>G358S</sup> | SJ733 | *Pf*ATP4<sup>G358S</sup> | 0.22 ± 0.018<br>(0.6)** | 1.0 ± 0.11<br>(1.9)** | 755 ± 79<br>(580)** |
| Dd2<sup>L350H</sup> | **15** | *Pf*ATP4<sup>L350H</sup> | 1.6 ± 0.16<br>(4.2)** | 3.4 ± 0.32<br>(6.3)* | 4.8 ± 0.30<br>(3.7)** |
| Dd2<sup>P412L</sup> | **15** | *Pf*ATP4<sup>P412L</sup> | 34 ± 1.3<br>(89)** | 4.1 ± 0.45<br>(7.6)* | 11 ± 0.70<br>(8.5)** |

EC<sub>50</sub> values are presented as means ± SEM from at least four independent experiments derived from duplicate technical replicates. Statistical significance of EC<sub>50</sub> fold-changes (in parentheses) was determined for all mutants against the Dd2-B2 parental line using two-tailed Mann-Whitney *U* tests (GraphPad Prism 10). \*, *P* < 0.05; \*\*, *P* < 0.01. Selection of the SJ733 resistant line (G358S) was previously reported<sup>27</sup>; selections using **15** yielded L350H and P412L (see supporting information). WT, wildtype; <sup>a</sup> Dd2-B2 is a clone of Dd2; WT Dd2 strains are resistant to chloroquine, mefloquine, and pyrimethamine.

## DISCUSSION

Malaria remains one of the most significant infectious diseases impacting populations globally. The capacity of *P. falciparum* to evade drug therapies through resistance presents significant challenges to treatment and control efforts. ^8–9^ This observation underlines the urgent necessity for identifying and advancing new antimalarial compounds into clinical development. Phenotypic HTS has emerged as a key method to discover new chemical classes for drug development against *P. falciparum.* ^9, 14^ Herein, we used a phenotypic screen to identify several α-azacyclic acetamide-based fast-killing antimalarials (**1**-**4**) with low micromolar activity against ABS parasites. These compounds demonstrate favorable drug-like properties including good solubility, and they were non-cytotoxic to the human HepG2 cells, making them candidates for further hit-to-lead optimization. We went on to show, using both biochemical assays and forward genetics, that the α-azacyclic acetamides target *Pf*ATP4 and have similar cell-based effects on *P. falciparum* as the clinical candidate cipargamin (KAE609), the first identified *Pf*ATP4 inhibitor to reach clinical development. ^20, 22–23^

To gain insight into the MOA of the α-azacyclic acetamides, we first sought to determine if they hit any known targets, ruling out DHODH and bc1 using a resistant transgenic parasite line. Previous studies have shown that the MOA of antimalarial compounds correlates with their kill rate.^36 34^ We therefore conducted biochemical assays to determine if the α-azacyclic acetamides might inhibit the plasma membrane ATP-dependent transporter *Pf*ATP4 because inhibitors of this target also show fast-killing kinetics.^24–25^ Using established cell-based measurements of pH and [Na^+^] ^41–44^ we found that similarly to KAE609, the α-azacyclic acetamides cause dose-dependent increases in parasite cytoplasmic pH and intracellular Na^+^ concentration. Building on prior assay approaches we showed that the EC_50_ for *P. falciparum* growth inhibition in the whole cell assay was approximately equivalent to the value derived from pH assays, providing strong evidence that the MOA of the α-azacyclic acetamides was inhibition of *Pf*ATP4. In addition to their effects on pH_cyt_ and [Na^+^]_cyt_, *Pf*ATP4 inhibitors give rise to swelling of both the parasite and parasitized erythrocyte, with the swelling contributing to parasite killing.^49^ Based on the common impact on intracellular pH and Na^+^, cell swelling is also likely to contribute to parasite killing by the α-azacyclic acetamides.

Resistance selections using diverse compounds that target *Pf*ATP4 have identified over 40 mutations in *Pf*ATP4, highlighting the genetic plasticity of *Pfatp4* and its ability to adapt to drug pressure.^19, 21, 27, 38, 50–53^ These selections have also proven to be a robust mechanism for identifying *Pf*ATP4 as the target for many of these compounds. A study assessing the *ex vivo* susceptibilities of clinical isolates in Uganda to KAE609 and SJ733 identified a number of polymorphisms in *Pfatp4* that led to modest decreases in compound efficacy, again highlighting the adaptability of the target.^54^ Here, further confirmation of the MOA of the α-azacyclic acetamides and insights into their binding modes was obtained by selection and profiling of **2** and **3** resistant parasites, leading to the discovery of two single point mutations within the *Pf*ATP4 gene, P412L and F917L, which conferred resistance to all four α-azacyclic acetamides. These mutations, which map to the transmembrane helices of *Pf*ATP4, have been previously documented in other studies following selections for resistance to SJ733 and KAE609, respectively.^27, 52^ Cross-resistance studies on these lines as well as three lines that were selected for resistance to SJ733 or its analog **15**, harboring either a L350H, G358S, or P412L mutation in *Pf*ATP4, suggested that the α-azacyclic acetamides bind to *Pf*ATP4 in a site that overlaps the binding site of KAE609 and SJ733, but that their binding sites also have distinct features. Both the P412L and F917L mutations caused resistance to all 4 α-azacyclic acetamides and to KAE609, though **1** and **4** were less impacted by the P412L mutation than the other compounds. Likewise, the L350H mutation identified in **15** selections led to a similar shift in EC_50_ for the α-azacyclic acetamides and for KAE609, consistent with overlapping binding modes. In contrast, the potency of the a-azacyclic acetamides was not greatly affected by the G358S mutation that was previously identified in selections using SJ733 and detected in clinical samples from patients treated with KAE609. ^21, 23, 27^ This mutation confers high-level resistance to KAE609 and the relative insensitivity of the α-azacyclic acetamides to this mutation suggests that their binding modes do not fully overlap. These data also suggest that if the G358S mutation were to become fixed in the population through use of KAE609 as a drug therapy, that the α-azacyclic acetamides could provide a starting point to develop a new clinical candidate that does not share this resistance risk. Importantly, none of the described natural allelic variations in the clinical isolates were identified in the resistance selections for the α-azacyclic acetamides.

## CONCLUSION

As the global threat of malaria continues to persist and escalates, addressing this public health challenge through the development of novel treatments and prophylactic agents has become increasingly urgent. Our research efforts identified four promising α-azacyclic acetamides, which exhibit rapid and potent activity against ABS *P. falciparum* parasites. We observed a significant correlation between the potency of compounds in inducing *Pf*ATP4-associated physiological effects, such as disruption of parasite cytosolic pH and Na^+^ homeostasis, and their efficacy in inhibiting parasite growth, suggesting that ionic dysregulation contributes to their antimalarial activity. Furthermore, our studies show that resistance to these novel compounds is associated with the *Pf*ATP4 target. Notably, the compounds we investigated are not impacted by the *Pf*ATP4^G358S^ mutation that was clinically associated with high-level resistance to KAE609.^23^ In summary, if current *Pf*ATP4 clinical candidates are found to be inadequate in combating malaria, the α-azacyclic acetamides we have described herein hold significant potential as future starting points for a lead optimization program aimed at identifying alternative ATP4 inhibitors.

## EXPERIMENTAL SECTION

### Materials

The 100K UT Southwestern compound collection was sourced from Chemical Div. and ChemBridge. SW260412 (SW412) (**1**) (*N*-(5-chloro-2-methoxyphenyl)-2-(2-(5-cyclopropyl-1,3,4-oxadiazol-2-yl)-1*H*-pyrrol-1-yl)acetamide, SW284491 (SW491) (**2**) (*N*-(4-methylpyridin-2-yl)-2-(2-(4-phenylthiazol-2-yl)-1*H*-pyrrol-1-yl)acetamide), SW262968 (SW968) (**3**) (*N*-(2-(4-chloro-3,5-dimethyl-1*H*-pyrazol-1-yl)ethyl)-2-(3-cyano-2-methyl-1*H*-indol-1-yl)acetamide) and SW302080 (SW080) (**4**) (*N*-(3-fluorophenyl)-2-(6-(5-methyl-1,3,4-oxadiazol-2-yl)indolin-1-yl)acetamide) were purchased from Sigma as were analogs shown in Table S1. KAE609 (Cat. No. 30678) was purchased from Cayman Chemical, artemisinin (HY-B0094) was purchased from Med Chem Express and concanamycin A was purchased from AdipoGen (BVT-0237). MMV609 (**15**)(compound 1a ^48^) was obtained from Medicines for Malaria Venture. DSM265 was synthesized as previously described.^17^

#### High throughput screen (HTS)

The α-azacyclic acetamides (**1**-**4**) were identified in a previously reported phenotypic screen for inhibitors of *P. falciparum* ABS parasite growth that was conducted as described. ^29–30^ However, the prior publications focused on different hits and the α-azacyclic acetamides were not previously disclosed.

#### *P. falciparum* asexual blood stage culture for EC_50_ determination (Phillips Lab)

ABS *P. falciparum* 3D7, Dd2 or *Pf*NF54^luc^ parasites (BEI resources) were cultured in RPMI 1640 medium (Millipore Sigma), supplemented with 25 mM HEPES, 0.5% Albumax-I (Thermo Scientific), 23 mM sodium bicarbonate, 92 µM hypoxanthine, and 12.5 µg/mL gentamicin sulfate at 37°C, 5% CO_2_, using deidentified male O^+^ red blood cells (BioIVT) as previously described. ^28–30^ Growth inhibition assays were conducted in 384-well plate format using the SYBR green method with modifications as described ^30^ (see supporting information for a detailed description).

#### *P. falciparum* asexual blood stage culture for cross-resistance profiling (Fidock Lab)

ABS parasites were cultured at 2% hematocrit in human O+ or A+ RBCs in RPMI-1640 media, supplemented with 25 mM HEPES, 50 mg/L hypoxanthine, 2 mM L-glutamine, 0.21% sodium bicarbonate, 0.5% (wt/vol) AlbuMAXII (Invitrogen) and 10 μg/mL gentamycin, in modular incubator chambers (Billups-Rothenberg) at 5% O_2_, 5% CO_2_ and 90% N_2_ at 37°C.

#### Human HepG2 cell culture and cytotoxicity assays

Cytotoxicity assays were conducted in HepG2 cells (ATCC) grown in EMEM medium supplemented with 10% fetal bovine serum, 1% Pen/Strep and 2 mM L-glutamine at 37°C in 5% CO_2_ and cell growth was monitored using a luciferase-coupled ATP quantification assay (Promega-CellTiter Glo®) as described.^30^

#### Data analysis and curve fitting

EC_50_ and CC_50_ values were determined by non-linear regression performed in GraphPad Prism Version 10. Unless otherwise specified, data were fitted to the log(inhibitor) vs response—variable slope with four parameters equation. In the displayed graphs, data were normalized to define 100% as the average value of the DMSO control and the 0% baseline was defined based on the highest dose used versus wild-type cells. For mutant cell lines and compound combinations where bottom plateau of zero growth was not reached, the bottom of the curve was defined at 0% based on background values obtained for parental Dd2 from the same study.

#### Rate of kill assay bioluminescence relative rate of kill (BRRoK) assay

Compound kill rate was evaluated using the BRRok assay performed on *P. falciparum* NF54^luc^ parasites (0.1mL, 4% hematocrit, 2% parasitemia) at the trophozoite-stage (20-26 h post infection) after a 6 h incubation with compounds over a concentration range (30, 18, 11, 6.7, 4.1, 2.5, 1.5, 0.92, 0.54, 0.33, and 0.10 ×EC₅₀) as described. ^28, 34^ Bioluminescence was used as a readout for viability at the end of the incubation (luciferase assay system; Promega) (see supporting information).

#### *P. falciparum* cytosolic dose response pH Assay

Cytosolic pH (pH_cyt_) effects were measured as described previously.^44^ Detailed methods are also provided in supporting information. Briefly, pH_cyt_ assays were carried out using *Pf*3D7 trophozoite-stage parasites (20-26h post infection) which were obtained via synchronization with a 5% w/v sorbitol. Saponin isolated parasites suspended in Albumax free RPMI medium pH 7.1, were loaded with the pH-sensitive fluorescent dye ester, BCECF-AM (Thermo Fisher Scientific B1170) and prepared as described in supporting informaiton. Prior to addition of compounds parasites were resuspended in glucose saline solution (125 mM NaCl, 25 mM HEPES, 5 mM KCl, 1 mM MgCl_2_, 20 mM glucose) pH 7.1 buffer, which is the estimated pH of the cytosol of the parasitized erythrocyte and plated onto 96-well plates at ∼10^8^ parasites per well. CMA (50 nM) was added first followed by test compounds once baseline fluorescence was established (final DMSO concentration 2%). Fluorescence was monitored at 37°C on a BioTek Synergy H1 Hybrid plate reader set for 440 nm and 490 nm excitation and 535 nm emission. Standard curves were also prepared using cells that were suspended in pH calibration buffer (130 mM KCl, 1 mM MgCl_2_, 20 mM glucose, 25 mM HEPES; pH 6.8, 7.1, 7.8).

The impact on cytosolic pH on *Pf*3D7 parasites was assessed over a range of compound concentrations (40 - 0.002 times the fold EC_50_ with an 8-point dilution series). pH data were averaged over a 10-minute period selected based on the time frame corresponding to the maximal pH change observed after CMA addition in the controls (DSM265 and CMA). Averaging the data over this period provided a robust and biologically relevant measure of the acidification effect for each experimental condition. To comprehensively evaluate compound efficacy, pH changes were plotted alongside *P. falciparum* growth inhibition data collected on independent cells. The average pH values (right-hand axis) over the defined 10-minute interval were compared to growth inhibition results (left-hand axis) across the same concentration range.

#### *P. falciparum* pH fingerprint assay

These studies were performed as previously described ^41^ and details are provided in supporting information.

#### Measurement of P. falciparum cytosolic Na^+^ concentrations

Cytosolic Na^+^ concentrations were measured in saponin-isolated trophozoite-stage *P. falciparum* parasites (3D7 strain) that were loaded with the Na^+^-sensitive fluorescent dye SBFI (Molecular Probes, Invitrogen, S1263) and suspended at 37°C in glucose-containing saline (125 mM NaCl, 5 mM KCl, 1 mM MgCl_2_, 20 mM glucose and 25 mM HEPES; pH 7.1) (as described previously). ^24^ Measurements were carried out in 96 well plates (excitation wavelengths: 340 nm and 380 nm; emission wavelength: 515 nm), as described previously.^37^ Fluorescence Ratio values (340 nm/380 nm) were converted to [Na^+^]_cyt_ using a calibration procedure in which parasites were suspended in solutions of varying [Na^+^] containing ionophores.^24^ Test and control compounds were diluted to their final concentrations in the assay from DMSO stocks, giving rise to a final DMSO concentration in the assay of 0.1% v/v.

#### Selection of 2- and 3- resistant parasites (Phillips Lab)

Resistant Dd2 parasites were selected using a high-pressure intermittent selection protocol using a previously described method.^11^ Selections were set up using 4 flasks per compound starting with approximately 3×10^9^ parasites per flask (80 mL per flask, 4% hematocrit, 3% parasitemia at the beginning of each pulse, mostly ring-stage parasites via previous sorbitol synchronization. Pulse 1 and 2 (48 h) were conducted at 10xEC_50_, and subsequent pulses were conducted at 20xEC_50_. Parasites cleared within 48 h under these conditions. **3**-selections used a total of 4 pulses, while **2**-selections required 5 pulses before resistant parasites emerged (Table S3). Culture media was replenished daily for a week after each pulse, then every two days with weekly replenishment of blood. Cultures were monitored by microscopy until recrudescence. Clonal parasites were obtained from each flask by limiting dilution and maintained in drug free RPMI-1640 media prior to being analyzed for EC_50_ shifts. Established clonal lines were aliquoted and stored at −80°C for future use after suspension in Glycerolyte 57 solution (Baxter Cat no 4A7831).

#### Statistical Analysis

Statistical analysis of EC_50_ shifts between the mutant and wild-type Dd2 cells was conducted as described in the table legends (Tables 2 and 3).

#### Parasite lines for cross resistance profiling and selection of MMV609 resistant lines (Fidock Lab)

Three *Pf*ATP4 mutant clones generated by *in vitro* selections of Dd2-B2 (a clone of Dd2) with the dihydroisoquinolone SJ733 (*Pf*ATP4^G358S^)^27^ or its analog MMV609 (*Pf*ATP4^L350H^ and *Pf*ATP4^P412L^) were assessed for susceptibility to **2**, **3** and KAE609. The Dd2 G358S mutant is a KAE609- and SJ733-resistant clone (DD2-SJ16-D2) that emerged following *in vitro* selection with SJ733.^27^ The L350H and P412L mutants were selected *in vitro* under 10× EC_50_ MMV609 pressure and exhibited 30-fold and 1850-fold EC_50_ increases against **15**, respectively, relative to Dd2 (see supporting information and Table S8 for details).

#### Sample preparation for whole genome sequencing (WGS)

Genomic DNA samples were prepared from four clones derived from **2**-selections (one deriving from each parent flask), three clones from **3**-selections (one deriving from 3 of 4 parent flasks), and three parental Dd2 clones. Infected RBCs suspended in PBS (pH 7.4) were subjected to 0.1% saponin treatment followed by centrifugation at 2,500xg with a brake speed of 50% to pellet the released parasites and remove host cell debris. Pellets were then washed three times with cold 1xPBS in a microcentrifuge tube, with centrifugation at 3,300xg after each wash. Parasites were then suspended in 200 µL PBS (pH 7.4) and genomic DNA was isolated from parasites using a Qiagen Blood and Cell Culture DNA Mini kit. Two changes were made to the manufacturer’s instructions: 1) 400 µg (4 µL at 100 mg/mL) of RNase A was added to 200 µL along with the recommended proteinase K; 2) samples were eluted in water instead of the provided buffer.

#### Whole-genome sequencing (WGS) and analysis

Paired-end sequencing was performed with 150-base reads using Illumina NextSeq 500. WGS data have been deposited in NCBI’s Sequence Read Archive (SRA) database (SRA BioProject ID PRJNA1230203). The sequenced reads per sample were subjected to analysis for quality and contamination using FastQC v0.11.260 and FastQ Screen v0.4.4.^55^ respectively. The reads from each sample were then mapped to the *Plasmodium falciparum* Dd2 reference genome using BWA-MEM,^56^ with default parameters. BWA efficiently aligns short-read sequences against a reference genome, allowing gaps and mismatches. Duplicated reads were marked, and coverage was calculated using Picard tools (v1.127). Potential SNPs and indels were discovered by running GATK’s (v3.5) HaplotypeCaller (parameters: -ploidy 1, --emitRefConfidence GVCF). Genotyping was then performed on samples pre-called with HaplotypeCaller using GenotypeGVCFs. A hard-filtering approach was used to filter variants. Briefly, the SelectVariants module was used to subset SNPs and INDELS. Then, the VariantFilteration module was used to filter variants. Variants were annotated using SnpEff.^57^ Gene and protein analysis for genes with coding changes was performed using PlasmoDB (release 68) for the following ID entries: *PfATP4* (PfDd2_120016700), *Pf*S8e (PfDd2_070012100), *Pf* Conserved protein (PfDd2_130068800) and *Pf*MDR1 (PfDd2_050027900).

#### Sanger sequencing of pfatp4, pfrs8e, pfcuf, and pfmdr1

To verify mutations identified via WGS, regions of interest were PCR-amplified from genomic DNA (isolated as stated for WGS) and submitted for sequencing at either the UT Southwestern Sanger Sequencing Core (Figures S6 and S7) or to Plasmidsaurus (Figures S8-S10), which performs primeless long reading sequencing. Primers for amplification and gene ID information used for analysis of the **2**- and **3**-resistant clones and matched parental Dd2 lines can be found in Table S6. The resulting sequencing analysis for these genes can be found in Figure S6-10. Sanger sequencing of *pfatp4* to identify mutations in the gene as the result of the **15**-selections were performed with primers described in Table S7.

#### Mapping of mutations onto an Alpha Fold generated homology model

Homology modeling was performed using AlphaFold (version 4), with structural predictions generated using the v2.3.0 model for the P-type ATPase Q27724. Models were visualized in PyMOL (Schrödinger, LLC, version 3.13).

#### In vitro ADME

Chemical properties (molecular weight, polar surface area, freely rotating bonds, hydrogen bond donors and acceptors, predicted pKa, and cLogD (pH 7.4)) were calculated using ChemAxon chemistry cartridge with JChem for Excel software (version 16.4.11). Solubility measurements and assessment of microsome stability was conducted as previously described.^17^

## ASSOCIATED CONTENT

### SUPPORTING INFORMATION

Additional supporting tables and figures, and detailed experimental methods, EC_50_ data for commercially sourced analogs, supporting data for resistance selections, supporting data for intracellular *P. falciparum* pH measurements, sequencing and PCR primers, supporting sequencing data (pdf).

## AUTHOR INFORMATION

### AUTHOR CONTRIBUTIONS

Conceptualization, M.A.P. (Biology), J.M.R. (Chemistry) and S.A.C. (Pharmacology); Methodology, A.C., L.S.I., V.T., A.M.L., A.K.L., S.L., A.K., C.X., H.N., B.A.P., B.L., S.A.C., D.A.F., J.M.R., and M.A.P.; Formal Analysis, A.C., L.S.I., V.T., A.K.L., A.M.L., S.A.C., D.A.F., J.M.R., and M.A.P.; Investigation, A.C., L.S.I., V.T., I.D., A.M.L, A.K.L., K.J.F., J.S., S.L., A.K., C.X., H.N., and; Resources, M.A.P., J.M.R., S.A.C. A.M.L. and D.A.F.; Data Curation, A.K. and C.X.,; Supervision, M.A.P., J.M.R., S.A.C., D.A.F. Writing – Original Draft, A.C., M.A.P. and J.M.R.; Writing – Review & Editing, All authors; Project Administration, M.A.P., J.M.R., S.A.C. and B.L.; Funding Acquisition, M.A.P., J.M.R., S.A.C., and D.A.F.

## ACKNOWLEDGEMENTS

This work was funded by the United States National Institutes of Health grants R01AI155784 (to MAP and JMR), R01AI103947 (to MAP), R01AI188859 (to DAF), T32GM007062 (to AKL), and the Medicines for Malaria Venture. SAC receives support from Monash University Technology Research Platform network and Therapeutic Innovation Australia (TIA) through the Australian Government National Collaborative Research Infrastructure Strategy (NCRIS) program. MAP and JMR acknowledge support of the Welch Foundation (I-1257, I-1612). BAP acknowledges the NIH-funded Shared Instrumentation Grant 1S10OD026758-01 for the Echo 655 used to add compounds in dose-response experiments with HepG2 cells. MAP holds the Sam G. Winstead and F. Andrew Bell Distinguished Chair in Biochemistry and JMR holds the Ronald W. Estabrook, Ph.D. Distinguished University Chair in Biomedical Science. The funders had no role in the study design, data collection and analysis, decision to publish, or preparation of the manuscript. The authors would like to thank Didier Leroy from Medicines for Malaria Venture for his contribution to the *Pf*ATP4 resistance studies. We are grateful to the Canberra Branch of the Australian Red Cross Lifeblood for the provision of blood. The authors would also like to acknowledge the PlasmoDB data base resource that facilitated studies described in this manuscript.

## ABBREVIATIONS

*Pf*ATP4: *P. falciparum* Na^+^ pump
KAE609: cipargamin
WHO: World Health Organization
ABS: asexual blood stage
HTS: high-throughput screening
ART: artemisinin
ACTs: artemisinin combination therapies
DHODH: dihydroorotate dehydrogenase
WGS: whole genome sequencing
HTS: high-throughput screen
HepG2: human hepatic cell line
yDHODH: *P. falciparum* strain expressing *Saccharomyces cerevisiae* yeast DHODH
ADME: Absorption, distribution, metabolism and excretion
SAR: structure activity relationship
BRRoK: bioluminescence relative rate of kill
MOA: mechanism of action
MMV: Medicines for Malaria Venture’s
[Na^+^]_cyt_: parasite’s cytosolic sodium concentration
(pH_cyt_): cytosolic pH
CMA: concanamycin A
EC_50_: effective concentration to reach 50%
RBC: red blood cell, DMSO, dimethyl sulfoxide
CL_int_: intrinsic clearance
CC_50_: cytotoxic concentration to reach 50%
*Pf*3D7: *P. falciparum* 3D7
*Pf*Dd2: *P. falciparum* Dd2
*Pf* NF54^luc^: *P. falciparum* transgenic strain expressing luciferase

---

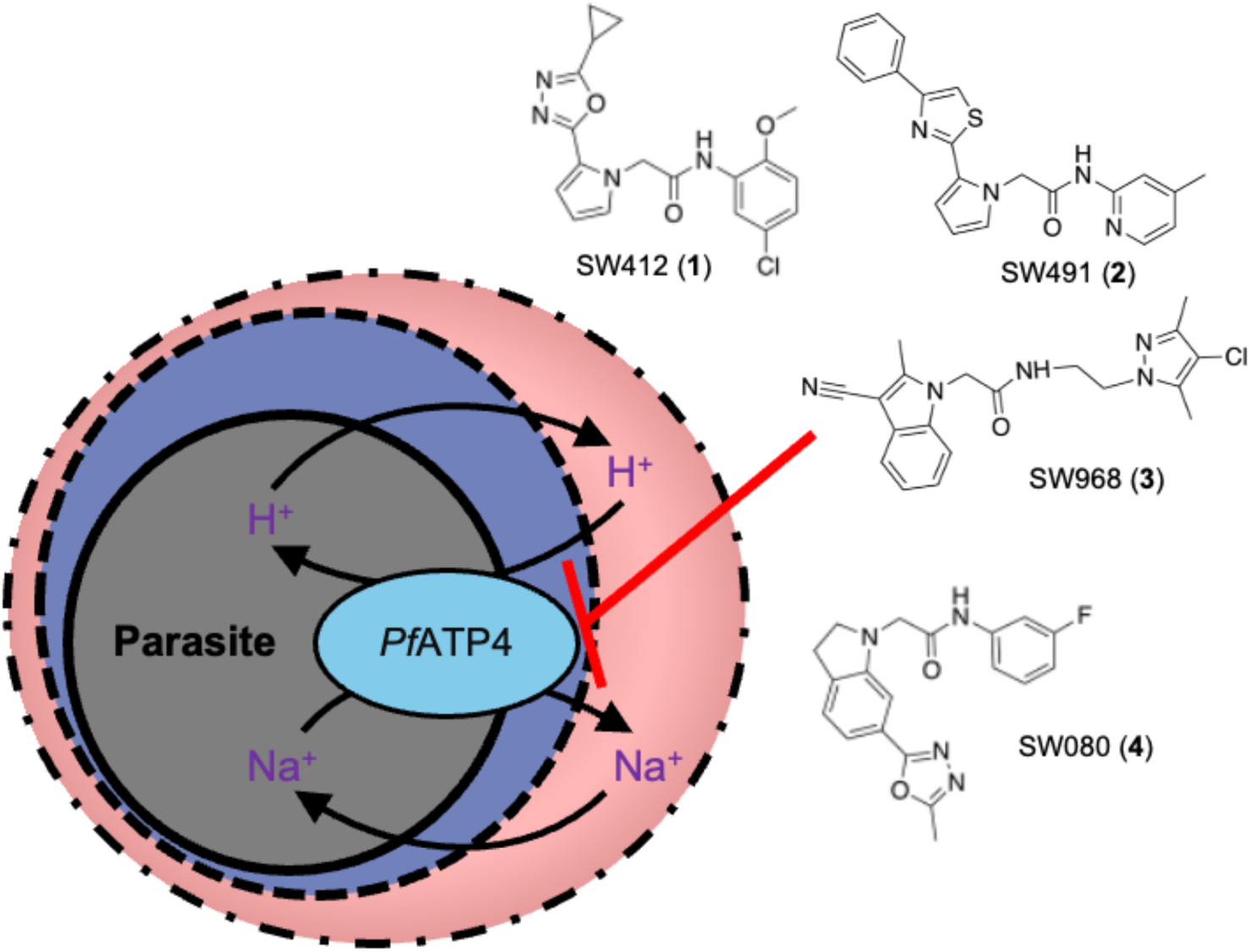

This TOC graphic shows the newly identified α-azacyclic acetamides have antimalarial activity in asexual blood-stage parasites and target *Pf*ATP4 (*P. falciparum* Na^+^ pump).

## REFERENCES

1. WHO World Malaria Report 2024. (accessed Feb 11 2025).

2. Mihreteab, S.; Platon, L.; Berhane, A.; Stokes, B. H.; Warsame, M.; Campagne, P.; Criscuolo, A.; Ma, L.; Petiot, N.; Doderer-Lang, C.; Legrand, E.; Ward, K. E.; Zehaie Kassahun, A.; Ringwald, P.; Fidock, D. A.; Menard, D., Increasing prevalence of artemisinin-resistant HRP2-negative malaria in Eritrea. N Engl J Med 2023, 389, 1191–1202.

3. Conrad, M. D.; Asua, V.; Garg, S.; Giesbrecht, D.; Niare, K.; Smith, S.; Namuganga, J. F.; Katairo, T.; Legac, J.; Crudale, R. M.; Tumwebaze, P. K.; Nsobya, S. L.; Cooper, R. A.; Kamya, M. R.; Dorsey, G.; Bailey, J. A.; Rosenthal, P. J., Evolution of partial resistance to artemisinins in malaria parasites in Uganda. N Engl J Med 2023, 389, 722–732.

4. Tilley, L.; Straimer, J.; Gnadig, N. F.; Ralph, S. A.; Fidock, D. A., Artemisinin action and resistance in Plasmodium falciparum. Trends Parasitol 2016, 32, 682–696.

5. Uwimana, A.; Legrand, E.; Stokes, B. H.; Ndikumana, J.-L. M.; Warsame, M.; Umulisa, N.; Ngamije, D.; Munyaneza, T.; Mazarati, J.-B.; Munguti, K.; Campagne, P.; Criscuolo, A.; Ariey, F.; Murindahabi, M.; Ringwald, P.; Fidock, D. A.; Mbituyumuremyi, A.; Menard, D., Emergence and clonal expansion of in vitro artemisinin-resistant *Plasmodium falciparum* kelch13 R561H mutant parasites in Rwanda. Nat Med 2020, 26, 1602–1608.

6. Poespoprodjo, J. R.; Douglas, N. M.; Ansong, D.; Kho, S.; Anstey, N. M., Malaria. Lancet 2023, 402, 2328–2345.

7. Moxon, C. A.; Gibbins, M. P.; McGuinness, D.; Milner, D. A.; Marti, M., New Insights into malaria pathogenesis. Annu Rev Pathol 2020, 15, 315–343.

8. Phillips, M. A.; Burrows, J. N.; Manyando, C.; van Huijsduijnen, R. H.; Van Voorhis, W. C.; Wells, T. N. C., Malaria. Nat Rev Dis Primers 2017, 3, 17050.

9. Siqueira-Neto, J. L.; Wicht, K. J.; Chibale, K.; Burrows, J. N.; Fidock, D. A.; Winzeler, E. A., Antimalarial drug discovery: progress and approaches. Nat Rev Drug Discov 2023, 22, 807–826.

10. Birkholtz, L.-M.; Alano, P.; Leroy, D., Transmission-blocking drugs for malaria elimination. Trends Parasitol 2022, 38, 390–403.

11. Corey, V. C.; Lukens, A. K.; Istvan, E. S.; Lee, M. C. S.; Franco, V.; Magistrado, P.; Coburn-Flynn, O.; Sakata-Kato, T.; Fuchs, O.; Gnädig, N. F.; Goldgof, G.; Linares, M.; Gomez-Lorenzo, M. G.; De Cózar, C.; Lafuente-Monasterio, M. J.; Prats, S.; Meister, S.; Tanaseichuk, O.; Wree, M.; Zhou, Y.; Willis, P. A.; Gamo, F.-J.; Goldberg, D. E.; Fidock, D. A.; Wirth, D. F.; Winzeler, E. A., A broad analysis of resistance development in the malaria parasite. Nat Commun 2016, 7, 1–9.

12. Burrows, J. N.; Duparc, S.; Gutteridge, W. E.; Hooft van Huijsduijnen, R.; Kaszubska, W.; Macintyre, F.; Mazzuri, S.; Mohrle, J. J.; Wells, T. N. C., New developments in anti-malarial target candidate and product profiles. Malar J 2017, 16, 26.

13. Katsuno, K.; Burrows, J. N.; Duncan, K.; van Huijsduijnen, R. H.; Kaneko, T.; Kita, K.; Mowbray, C. E.; Schmatz, D.; Warner, P.; Slingsby, B. T., Hit and lead criteria in drug discovery for infectious diseases of the developing world. Nat Rev Drug Discov 2015, 14, 751–758.

14. Hovlid, M. L.; Winzeler, E. A., Phenotypic screens in antimalarial drug discovery. Trends Parasitol 2016, 32, 697–707.

15. Antonova-Koch, Y.; Meister, S.; Abraham, M.; Luth, M. R.; Ottilie, S.; Lukens, A. K.; Sakata-Kato, T.; Vanaerschot, M.; Owen, E.; Jado, J. C.; Maher, S. P.; Calla, J.; Plouffe, D.; Zhong, Y.; Chen, K.; Chaumeau, V.; Conway, A. J.; McNamara, C. W.; Ibanez, M.; Gagaring, K.; Serrano, F. N.; Eribez, K.; Taggard, C. M.; Cheung, A. L.; Lincoln, C.; Ambachew, B.; Rouillier, M.; Siegel, D.; Nosten, F.; Kyle, D. E.; Gamo, F. J.; Zhou, Y.; Llinas, M.; Fidock, D. A.; Wirth, D. F.; Burrows, J.; Campo, B.; Winzeler, E. A., Open-source discovery of chemical leads for next-generation chemoprotective antimalarials. Science 2018, 362, eaat9446.

16. Delves, M. J.; Miguel-Blanco, C.; Matthews, H.; Molina, I.; Ruecker, A.; Yahiya, S.; Straschil, U.; Abraham, M.; Leon, M. L.; Fischer, O. J.; Rueda-Zubiaurre, A.; Brandt, J. R.; Cortes, A.; Barnard, A.; Fuchter, M. J.; Calderon, F.; Winzeler, E. A.; Sinden, R. E.; Herreros, E.; Gamo, F. J.; Baum, J., A high throughput screen for next-generation leads targeting malaria parasite transmission. Nat Commun 2018, 9, 3805.

17. Phillips, M. A.; Lotharius, J.; Marsh, K.; White, J.; Dayan, A.; White, K. L.; Njoroge, J. W.; El Mazouni, F.; Lao, Y.; Kokkonda, S.; Tomchick, D. R.; Deng, X.; Laird, T.; Bhatia, S. N.; March, S.; Ng, C. L.; Fidock, D. A.; Wittlin, S.; Lafuente-Monasterio, M.; Benito, F. J.; Alonso, L. M.; Martinez, M. S.; Jimenez-Diaz, M. B.; Bazaga, S. F.; Angulo-Barturen, I.; Haselden, J. N.; Louttit, J.; Cui, Y.; Sridhar, A.; Zeeman, A. M.; Kocken, C.; Sauerwein, R.; Dechering, K.; Avery, V. M.; Duffy, S.; Delves, M.; Sinden, R.; Ruecker, A.; Wickham, K. S.; Rochford, R.; Gahagen, J.; Iyer, L.; Riccio, E.; Mirsalis, J.; Bathhurst, I.; Rueckle, T.; Ding, X.; Campo, B.; Leroy, D.; Rogers, M. J.; Rathod, P. K.; Burrows, J. N.; Charman, S. A., A long-duration dihydroorotate dehydrogenase inhibitor (DSM265) for prevention and treatment of malaria. Sci Transl Med 2015, 7, 296ra111.

18. Carolino, K.; Winzeler, E. A., The antimalarial resistome - finding new drug targets and their modes of action. Curr Opin Microbiol 2020, 57, 49–55.

19. Rottmann, M.; McNamara, C.; Yeung, B. K. S.; Lee, M. C. S.; Zou, B.; Russell, B.; Seitz, P.; Plouffe, D. M.; Dharia, N. V.; Tan, J.; Cohen, S. B.; Spencer, K. R.; González-Páez, G. E.; Lakshminarayana, S. B.; Goh, A.; Suwanarusk, R.; Jegla, T.; Schmitt, E. K.; Beck, H.-P.; Brun, R.; Nosten, F.; Renia, L.; Dartois, V.; Keller, T. H.; Fidock, D. A.; Winzeler, E. A.; Diagana, T. T., Spiroindolones, a potent compound class for the treatment of malaria. Science 2010, 329, 1175–1180.

20. Ndayisaba, G.; Yeka, A.; Asante, K. P.; Grobusch, M. P.; Karita, E.; Mugerwa, H.; Asiimwe, S.; Oduro, A.; Fofana, B.; Doumbia, S.; Jain, J. P.; Barsainya, S.; Kullak-Ublick, G. A.; Su, G.; Schmitt, E. K.; Csermak, K.; Gandhi, P.; Hughes, D., Hepatic safety and tolerability of cipargamin (KAE609), in adult patients with *Plasmodium falciparum* malaria: a randomized, phase II, controlled, dose-escalation trial in sub-Saharan Africa. Malar J 2021, 20, 478.

21. Qiu, D.; Pei, J. V.; Rosling, J. E. O.; Thathy, V.; Li, D.; Xue, Y.; Tanner, J. D.; Penington, J. S.; Aw, Y. T. V.; Aw, J. Y. H.; Xu, G.; Tripathi, A. K.; Gnadig, N. F.; Yeo, T.; Fairhurst, K. J.; Stokes, B. H.; Murithi, J. M.; Kumpornsin, K.; Hasemer, H.; Dennis, A. S. M.; Ridgway, M. C.; Schmitt, E. K.; Straimer, J.; Papenfuss, A. T.; Lee, M. C. S.; Corry, B.; Sinnis, P.; Fidock, D. A.; van Dooren, G. G.; Kirk, K.; Lehane, A. M., A G358S mutation in the *Plasmodium falciparum* Na(+) pump PfATP4 confers clinically-relevant resistance to cipargamin. Nat Commun 2022, 13, 5746.

22. Venishetty, V. K.; Lecot, J.; Nguyen, A.; Zhang, J.; Prince, W. T., First-in-human, randomized, double-blind, placebo-controlled, single and multiple ascending doses clinical study to assess the safety, tolerability, and pharmacokinetics of cipargamin administered intravenously in healthy adults. Antimicrob Agents Chemother 2024, 68, e0128723.

23. Schmitt, E. K.; Ndayisaba, G.; Yeka, A.; Asante, K. P.; Grobusch, M. P.; Karita, E.; Mugerwa, H.; Asiimwe, S.; Oduro, A.; Fofana, B.; Doumbia, S.; Su, G.; Csermak Renner, K.; Venishetty, V. K.; Sayyed, S.; Straimer, J.; Demin, I.; Barsainya, S.; Boulton, C.; Gandhi, P., Efficacy of Cipargamin (KAE609) in a randomized, phase II dose-escalation study in adults in Sub-Saharan Africa with uncomplicated *Plasmodium falciparum* malaria. Clin Infect Dis 2022, 74, 1831–1839.

24. Spillman, N. J.; Allen, R. J.; McNamara, C. W.; Yeung, B. K.; Winzeler, E. A.; Diagana, T. T.; Kirk, K., Na(+) regulation in the malaria parasite *Plasmodium falciparum* involves the cation ATPase PfATP4 and is a target of the spiroindolone antimalarials. Cell Host Microbe 2013, 13, 227–37.

25. Spillman, N. J.; Kirk, K., The malaria parasite cation ATPase PfATP4 and its role in the mechanism of action of a new arsenal of antimalarial drugs. Int J Parasitol Drugs Drug Resist 2015, 5, 149–62.

26. Gaur, A. H.; Panetta, J. C.; Smith, A. M.; Dallas, R. H.; Freeman, B. B., 3rd; Stewart, T. B.; Tang, L.; John, E.; Branum, K. C.; Patel, N. D.; Ost, S.; Heine, R. N.; Richardson, J. L.; Hammill, J. T.; Bebrevska, L.; Gusovsky, F.; Maki, N.; Yanagi, T.; Flynn, P. M.; McCarthy, J. S.; Chalon, S.; Guy, R. K., Combining SJ733, an oral ATP4 inhibitor of *Plasmodium falciparum*, with the pharmacokinetic enhancer cobicistat: An innovative approach in antimalarial drug development. EBioMedicine 2022, 80, 104065.

27. Jimenez-Diaz, M. B.; Ebert, D.; Salinas, Y.; Pradhan, A.; Lehane, A. M.; Myrand-Lapierre, M. E.; O’Loughlin, K. G.; Shackleford, D. M.; Justino de Almeida, M.; Carrillo, A. K.; Clark, J. A.; Dennis, A. S.; Diep, J.; Deng, X.; Duffy, S.; Endsley, A. N.; Fedewa, G.; Guiguemde, W. A.; Gomez, M. G.; Holbrook, G.; Horst, J.; Kim, C. C.; Liu, J.; Lee, M. C.; Matheny, A.; Martinez, M. S.; Miller, G.; Rodriguez-Alejandre, A.; Sanz, L.; Sigal, M.; Spillman, N. J.; Stein, P. D.; Wang, Z.; Zhu, F.; Waterson, D.; Knapp, S.; Shelat, A.; Avery, V. M.; Fidock, D. A.; Gamo, F. J.; Charman, S. A.; Mirsalis, J. C.; Ma, H.; Ferrer, S.; Kirk, K.; Angulo-Barturen, I.; Kyle, D. E.; DeRisi, J. L.; Floyd, D. M.; Guy, R. K., (+)-SJ733, a clinical candidate for malaria that acts through ATP4 to induce rapid host-mediated clearance of Plasmodium. Proc Natl Acad Sci U S A 2014, 111, E5455–62.

28. Lawong, A.; Gahalawat, S.; Okombo, J.; Striepen, J.; Yeo, T.; Mok, S.; Deni, I.; Bridgford, J. L.; Niederstrasser, H.; Zhou, A.; Posner, B.; Wittlin, S.; Gamo, F. J.; Crespo, B.; Churchyard, A.; Baum, J.; Mittal, N.; Winzeler, E.; Laleu, B.; Palmer, M. J.; Charman, S. A.; Fidock, D. A.; Ready, J. M.; Phillips, M. A., Novel antimalarial tetrazoles and amides active against the hemoglobin degradation pathway in *Plasmodium falciparum*. J Med Chem 2021, 64, 2739–2761.

29. Imlay, L. S.; Lawong, A. K.; Gahalawat, S.; Kumar, A.; Xing, C.; Mittal, N.; Wittlin, S.; Churchyard, A.; Niederstrasser, H.; Crespo-Fernandez, B.; Posner, B. A.; Gamo, F. J.; Baum, J.; Winzeler, E. A.; Laleu, B.; Ready, J. M.; Phillips, M. A., Fast-killing tyrosine amide ((s)-sw228703) with blood- and liver-stage antimalarial activity associated with the cyclic amine resistance locus (PfCARL). ACS Infect Dis 2023, 9, 527–539.

30. Lawong, A.; Gahalawat, S.; Ray, S.; Ho, N.; Han, Y.; Ward, K. E.; Deng, X.; Chen, Z.; Kumar, A.; Xing, C.; Hosangadi, V.; Fairhurst, K. J.; Tashiro, K.; Liszczak, G.; Shackleford, D. M.; Katneni, K.; Chen, G.; Saunders, J.; Crighton, E.; Casas, A.; Robinson, J. J.; Imlay, L. S.; Zhang, X.; Lemoff, A.; Zhao, Z.; Angulo-Barturen, I.; Jimenez-Diaz, M. B.; Wittlin, S.; Campbell, S. F.; Fidock, D. A.; Laleu, B.; Charman, S. A.; Ready, J. M.; Phillips, M. A., Identification of potent and reversible piperidine carboxamides that are species-selective orally active proteasome inhibitors to treat malaria. Cell Chem Biol 2024, 31, 1503–1517 e19.

31. Ganesan, S. M.; Morrisey, J. M.; Ke, H.; Painter, H. J.; Laroiya, K.; Phillips, M. A.; Rathod, P. K.; Mather, M. W.; Vaidya, A. B., Yeast dihydroorotate dehydrogenase as a new selectable marker for *Plasmodium falciparum* transfection. Mol Biochem Parasitol 2011, 177, 29–34.

32. Painter, H. J.; Morrisey, J. M.; Mather, M. W.; Vaidya, A. B., Specific role of mitochondrial electron transport in blood-stage *Plasmodium falciparum*. Nature 2007, 446, 88–91.

33. Lipinski, C. A.; Lombardo, F.; Dominy, B. W.; Feeney, P. J., Experimental and computational approaches to estimate solubility and permeability in drug discovery and development settings. Adv Drug Deliv Rev 2001, 46, 3–26.

34. Ullah, I.; Sharma, R.; Biagini, G. A.; Horrocks, P., A validated bioluminescence-based assay for the rapid determination of the initial rate of kill for discovery antimalarials. J Antimicrob Chemother 2017, 72, 717–726.

35. Sanz, L. M.; Crespo, B.; De-Cózar, C.; Ding, X. C.; Llergo, J. L.; Burrows, J. N.; García-Bustos, J. F.; Gamo, F.-J., *P. falciparum in vitro* killing rates allow to discriminate between different antimalarial mode-of-action. PloS one 2012, 7, e30949.

36. Burrows, J. N.; van Huijsduijnen, R. H.; Mohrle, J. J.; Oeuvray, C.; Wells, T. N., Designing the next generation of medicines for malaria control and eradication. Malar J 2013, 12, 187.

37. Dennis, A. S. M.; Rosling, J. E. O.; Lehane, A. M.; Kirk, K., Diverse antimalarials from whole-cell phenotypic screens disrupt malaria parasite ion and volume homeostasis. Sci Rep 2018, 8, 8795.

38. Lehane, A. M.; Ridgway, M. C.; Baker, E.; Kirk, K., Diverse chemotypes disrupt ion homeostasis in the Malaria parasite. Mol Microbiol 2014, 94, 327–39.

39. Rosling, J. E. O.; Ridgway, M. C.; Summers, R. L.; Kirk, K.; Lehane, A. M., Biochemical characterization and chemical inhibition of PfATP4-associated Na(+)-ATPase activity in *Plasmodium falciparum* membranes. J Biol Chem 2018, 293, 13327–13337.

40. Spillman, N. J.; Allen, R. J.; Kirk, K., Na+ extrusion imposes an acid load on the intraerythrocytic malaria parasite. Mol Biochem Parasitol 2013, 189, 1–4.

41. Lindblom, J. C. R.; Zhang, X.; Lehane, A. M., A pH Fingerprint Assay to Identify Inhibitors of Multiple Validated and Potential Antimalarial Drug Targets. ACS Infect Dis 2024, 10, 1185–1200.

42. Huss, M.; Ingenhorst, G.; Konig, S.; Gassel, M.; Drose, S.; Zeeck, A.; Altendorf, K.; Wieczorek, H., Concanamycin A, the specific inhibitor of V-ATPases, binds to the V(o) subunit c. J Biol Chem 2002, 277, 40544–8.

43. Kuhn, Y.; Rohrbach, P.; Lanzer, M., Quantitative pH measurements in *Plasmodium falciparum*-infected erythrocytes using pHluorin. Cell Microbiol 2007, 9, 1004–13.

44. Saliba, K. J.; Kirk, K., pH regulation in the intracellular malaria parasite, *Plasmodium falciparum*. H(+) extrusion via a V-type H(+)-ATPase. J Biol Chem 1999, 274, 33213–9.

45. Kirk, K., Ion Regulation in the Malaria Parasite. Annu Rev Microbiol 2015, 69, 341–59.

46. Das, S.; Bhatanagar, S.; Morrisey, J. M.; Daly, T. M.; Burns, J. M., Jr.; Coppens, I.; Vaidya, A. B., Na+ influx induced by new antimalarials causes rapid alterations in the cholesterol content and morphology of *Plasmodium falciparum*. PLoS Pathog 2016, 12, e1005647.

47. Cowell, A. N.; Istvan, E. S.; Lukens, A. K.; Gomez-Lorenzo, M. G.; Vanaerschot, M.; Sakata-Kato, T.; Flannery, E. L.; Magistrado, P.; Owen, E.; Abraham, M.; LaMonte, G.; Painter, H. J.; Williams, R. M.; Franco, V.; Linares, M.; Arriaga, I.; Bopp, S.; Corey, V. C.; Gnadig, N. F.; Coburn-Flynn, O.; Reimer, C.; Gupta, P.; Murithi, J. M.; Moura, P. A.; Fuchs, O.; Sasaki, E.; Kim, S. W.; Teng, C. H.; Wang, L. T.; Akidil, A.; Adjalley, S.; Willis, P. A.; Siegel, D.; Tanaseichuk, O.; Zhong, Y.; Zhou, Y.; Llinas, M.; Ottilie, S.; Gamo, F. J.; Lee, M. C. S.; Goldberg, D. E.; Fidock, D. A.; Wirth, D. F.; Winzeler, E. A., Mapping the malaria parasite druggable genome by using in vitro evolution and chemogenomics. Science 2018, 359, 191–199.

48. Guy, R. K.; Hammill, J. T.; Floyd, D.; Burrows, J.; Brand, S. New Anti-malarial agents. US 20230174503 AI, 2023.

49. Dennis, A. S. M.; Lehane, A. M.; Ridgway, M. C.; Holleran, J. P.; Kirk, K., Cell swelling induced by the antimalarial KAE609 (Cipargamin) and Other PfATP4-Associated Antimalarials. Antimicrob Agents Chemother 2018, 62.

50. Flannery, E. L.; McNamara, C. W.; Kim, S. W.; Kato, T. S.; Li, F.; Teng, C. H.; Gagaring, K.; Manary, M. J.; Barboa, R.; Meister, S.; Kuhen, K.; Vinetz, J. M.; Chatterjee, A. K.; Winzeler, E. A., Mutations in the P-type cation-transporter ATPase 4, PfATP4, mediate resistance to both aminopyrazole and spiroindolone antimalarials. ACS Chem Biol 2015, 10, 413–20.

51. Vaidya, A. B.; Morrisey, J. M.; Zhang, Z.; Das, S.; Daly, T. M.; Otto, T. D.; Spillman, N. J.; Wyvratt, M.; Siegl, P.; Marfurt, J.; Wirjanata, G.; Sebayang, B. F.; Price, R. N.; Chatterjee, A.; Nagle, A.; Stasiak, M.; Charman, S. A.; Angulo-Barturen, I.; Ferrer, S.; Belen Jimenez-Diaz, M.; Martinez, M. S.; Gamo, F. J.; Avery, V. M.; Ruecker, A.; Delves, M.; Kirk, K.; Berriman, M.; Kortagere, S.; Burrows, J.; Fan, E.; Bergman, L. W., Pyrazoleamide compounds are potent antimalarials that target Na+ homeostasis in intraerythrocytic *Plasmodium falciparum*. Nat Commun 2014, 5, 5521.

52. Lee, A. H.; Fidock, D. A., Evidence of a mild mutator phenotype in cambodian *Plasmodium falciparum* malaria parasites. PLoS One 2016, 11, e0154166.

53. Luth, M. R.; Godinez-Macias, K. P.; Chen, D.; Okombo, J.; Thathy, V.; Cheng, X.; Daggupati, S.; Davies, H.; Dhingra, S. K.; Economy, J. M.; Edgar, R. C. S.; Gomez-Lorenzo, M. G.; Istvan, E. S.; Jado, J. C.; LaMonte, G. M.; Melillo, B.; Mok, S.; Narwal, S. K.; Ndiaye, T.; Ottilie, S.; Palomo Diaz, S.; Park, H.; Pena, S.; Rocamora, F.; Sakata-Kato, T.; Small-Saunders, J. L.; Summers, R. L.; Tumwebaze, P. K.; Vanaerschot, M.; Xia, G.; Yeo, T.; You, A.; Gamo, F. J.; Goldberg, D. E.; Lee, M. C. S.; McNamara, C. W.; Ndiaye, D.; Rosenthal, P. J.; Schreiber, S. L.; Serra, G.; De Siqueira-Neto, J. L.; Skinner-Adams, T. S.; Uhlemann, A. C.; Kato, N.; Lukens, A. K.; Wirth, D. F.; Fidock, D. A.; Winzeler, E. A., Systematic *in vitro* evolution in *Plasmodium falciparum* reveals key determinants of drug resistance. Science 2024, 386, eadk9893.

54. Kreutzfeld, O.; Rasmussen, S. A.; Ramanathan, A. A.; Tumwebaze, P. K.; Byaruhanga, O.; Katairo, T.; Asua, V.; Okitwi, M.; Orena, S.; Legac, J.; Conrad, M. D.; Nsobya, S. L.; Aydemir, O.; Bailey, J.; Duffey, M.; Bayles, B. R.; Vaidya, A. B.; Cooper, R. A.; Rosenthal, P. J., Associations between Varied Susceptibilities to PfATP4 Inhibitors and Genotypes in Ugandan *Plasmodium falciparum* Isolates. Antimicrob Agents Chemother 2021, 65, e0077121.

55. Wingett, S. W.; Andrews, S., FastQ Screen: A tool for multi-genome mapping and quality control F1000Research 2018, 7, 1338.

56. Li, H., Aligning sequence reads, clone sequences and assembly contigs with BWA-MEM. arXiv 2013, 1303.3997v2, 1–3.

57. Cingolani, P.; Platts, A.; Wang, L. L.; Coon, M.; Nguyen, T.; Wang, L.; Land, S. J.; Lu, X.; Ruden, D. M., A program for annotating and predicting the effects of single nucleotide polymorphisms, SnpEff: SNPs in the genome of Drosophila melanogaster strain w1118; iso-2; iso-3. Fly 2012, 6, 80–92.

