## Supplemental methods, tables and figures for "Identification of α-azacyclic acetamide-based inhibitors of *P. falciparum* Na^+^ pump (*Pf*ATP4) with fast-killing asexual blood-stage antimalarial activity by phenotypic screening"

**Table of contents**


- **Supporting materials and methods........................................................................3-10**
- **Supporting tables...........................................................................................11-21**
  - **Table S1.** *P. falciparum* $EC_{50}$ data for commercially sourced analogs of **2** and **3**.
  - **Table S2.** Summary of *Pf*3D7 cellular and pH $EC_{50}$ data for DSM265 and KAE609
  - **Table S3.** Resistance Selection Cycle Recrudescence and Bulk culture $EC_{50}$ Data
  - **Table S4.** Protein coding mutations observed in **2**- and **3**- resistant parasites
  - **Table S5:** Cross resistance data for additional parasite clones resistant to **2** and **3**
  - **Table S6:** Primers for PCR amplification and sequencing Phillips Lab
  - **Table S7:** Primers used for PCR amplification and sequencing Fidock Lab
  - **Table S8:** Characteristics of Dd2 *Pf*ATP4 mutant lines used for cross-resistance profiling in Table 3
- **Supporting figures.........................................................................................21-27**
  - **Figure. S1.** α-azacyclic acetamides kill rate data, supporting data for Figure 1
  - **Figure S2.** The effects of **1**-**3** on intracellular pH, supporting data for Figure 2
  - **Figure S3.** pH versus time profiles for all intracellular pH, supporting data for Figure 2
  - **Figure S4.** pH versus time profiles for all intracellular pH, supporting data for Figure S2
  - **Figure S5.** *Pf*ATP4 pH fingerprint assay versus α-azacyclic acetamides **2** and **3**
  - **Figure S6.** Sanger sequencing to verify **2**-selected mutations in *pfatp4*
  - **Figure S7.** Sanger sequencing to verify **3**-selected mutations in *pfatp4*
  - **Figure S8.** Long read sequencing to verify **2**-selected mutations in Ribosomal protein S8e
  - **Figure S9.** Long read sequencing to verify **3**-selected mutations in *pfcpu*
  - **Figure S10.** Long read sequencing of multidrug resistance protein 1 *pfmdr1*

**Supplemental Materials.**

Analogs **5** (SW284463;SW463) N-(5-methylpyridin-2-yl)-2-[2-(4-phenyl-1,3-thiazol-2-yl)-1H-pyrrol-1-yl]acetamide, **6** (SW393316;SW316) N-(6-methylpyridin-2-yl)-2-[2-(4-phenyl-1,3-thiazol-2-yl)-1H-pyrrol-1-yl]acetamide, **7** (SW393317; SW317) 2-(2-(4-(4-chlorophenyl)-1,3-thiazol-2-yl)-1H-pyrrol-1-yl}-N-(4-methylpyridin-2-yl)acetamide, **8** (SW393318; SW318) 2-(2-(4-(4-chlorophenyl)-1,3-thiazol-2-yl)-1H-pyrrol-1-yl)-N-(3-methylpyridin-2-yl)acetamide, **9** (SW393319 ; SW319) N-(3-methylphenyl)-2-[2-(4-phenyl-1,3-thiazol-2-yl)-1H-pyrrol-1-yl]acetamide, **10** (SW393320; SW320) N-(3-chlorophenyl)-2-[2-(4-phenyl-1,3-thiazol-2-yl)-1H-pyrrol-1-yl]acetamide, **11** (SW214966; SW966) 2-(2-(4-(4-chlorophenyl)-1,3-thiazol-2-yl)-1H-pyrrol-1-yl)-1-(2,3-dihydro-1H-indol-1-yl)ethan-1-one, **12** (SW263181; SW181) 2-(3-acetyl-1H-indol-1-yl)-N-[2-(4-chloro-3,5-dimethyl-1H-pyrazol-1-yl)ethyl]propenamide, **13** (SW393314; SW314) 2-(1H-indol-1-yl)-N-[2-(1H-pyrazol-1-yl)ethyl]acetamide, **14** (SW393315; SW315) 2-(5-amino-3-methyl-1H-pyrazol-1-yl)-N-[2-(2-methyl-1H-indol-1-yl)ethyl]acetamide were purchased from Sigma.

**Supplemental Experimental Methods.**

***P. falciparum*** **growth inhibition assays to determine compound $EC_{50}$ (Phillips Lab).** Compounds were plated from 10 mM DMSO stocks using a Tecan D300e liquid handler to a final DMSO concentration of 0.5% over a range of concentrations in technical triplicate. Ring-stage parasites (200 μL/well) (2% hematocrit, 0.5% parasitemia) were added to 96-well black-walled clear flat-bottom plates and incubated at 37°C, 5% $CO_2$ for 72 h. Plates were then frozen (-80°C) overnight and then 100 µL of a working stock of SYBR® Green (Sigma- SYBR® Green nucleic acid gel stock) diluted 5000-fold in buffer (20 mM Tris-HCl pH 7.5, 5 mM EDTA, 0.008% w/vol saponin, 0.2% v/v Triton X-100, 0.0002% SYBR® Green) was added to each well and mixed thoroughly via pipetting. Fluorescence signal was read immediately on a BioTek Synergy H1 Hybrid plate reader set for 485 nm excitation and 535 nm emission. DSM265 was plated as a positive control. Statistical significance when comparing drug sensitivity between wild-type and resistant parasites was evaluated by ordinary one-way ANOVA analysis with Dunnett correction was performed in Graphpad Prism 10.

**Drug susceptibility assays for cross-resistance profiling (Fidock Lab).** To define the 50% ($EC_{50}$) and 90% ($EC_{90}$) growth-inhibitory concentrations for inhibitors **2**, **3**, and KAE609 for *Pf*ATP4 WT (Dd2-B2 parent) and G358S, L350H, and P412L mutant parasites, asynchronous cultures at 0.3-0.5% parasitemia and 1% hematocrit were exposed for 72 h to a range of ten compound concentrations that were two-fold serially diluted in duplicates along with compound-free controls. Parasite survival was assessed by flow cytometry on an Intellicyt iQue Screener PLUS (Sartorius) using 1× SYBR Green (Invitrogen) and 100 nM MitoTracker Deep Red FM (Invitrogen) as nuclear stain and vital dyes, respectively. $EC_{50}$ values were calculated from growth inhibition data using linear interpolation as means ± SEM from four to six independent experiments, each with technical duplicates. Statistical significance was determined against the Dd2-B2 parental control using two-tailed Mann-Whitney *U* tests (GraphPad Prism, version 10).

***In vitro* selection and characterization of 15-resistant *PfATP4* mutants (Fidock lab).** To generate mutants resistant to **15** (MMV1793609; MMV609), the B2 clone of the *P. falciparum* Dd2 strain was expanded to triplicate flasks each inoculated with 3E8 infected red blood cells (4% hematocrit). Parasites were exposed to constant **15** pressure at 10×$EC_{50}$ of the Dd2-B2 parent. Resistant parasites were detected in all three flasks after 12-17 days.

*Pfatp4* (PF3D7_1211900) was PCR amplified using KAPA HiFi HotStart ReadyMix (Roche) from genomic DNA extracted from **15**-resistant clones (obtained by limiting dilution) and the Dd2-B2 parent line. Amplification was performed using outer primers flanking the entire 3.8 kb gene: forward primer (p6536) 5'- ATGAGTTCTCAAAATAATAATAAACAGGGTGGAC and reverse primer (p6537) 5'- TTAATTCTTAATAGTCATATATTTTCTTCTATATATAACCTTTGG. PCR conditions were as follows: 95°C for 3 minutes, 45 rounds of 98°C for 20 seconds, 50°C for 30 seconds, and 68°C for 3 minutes, with a final extension of 4 minutes at 68°C. Agarose gel electrophoresis was used to confirm PCR product size; 15 sequencing of PCR products was carried out by Genewiz Inc. using the flanking and internal primers to obtain high quality, double-stranded sequence coverage (Table S7). Sequences were aligned to WT *pfatp4* (PF3D7_1211900) from the 3D7 genome reference strain (PlasmoDB, version 47) and analyzed on Geneious 9.1.8. Electropherograms were visually inspected to confirm the presence of *pfatp4* mutations and determine their allele frequencies.

Clones expressing the *Pf*ATP4 L350H and P412L mutations were identified and assessed for susceptibility to **15** and KAE609 (Table 3 and Table S8). In parallel, the previously reported SJ733-selected Dd2 G358S mutant clone (DD2-SJ16-D2) was tested against SJ733 and KAE609 to confirm the level of resistance conferred by this mutation (Table S8).

**Human HepG2 cell culture and cytotoxicity assays extended methods.** Cell growth was monitored using a luciferase-coupled ATP quantification assay (Promega-CellTiter Glo®) following manufacturer's instructions in 384-well white-walled opaque flat-bottom plate format. Cells were seeded to a density of 900 cells/well from a suspension ($1.5\times10^4$ cells/mL) and incubated overnight. Compounds were added the next day from DMSO stocks using a Labcyte Echo 655 acoustic dispenser to a final 0.5% DMSO concentration. Following treatment, cells were incubated at 37°C, 5% $CO_2$ for 96 h before addition of the CTG reagent and read out on a Revvity EnVision multimode plate reader. Assays were performed in technical triplicates. Brefeldin A was plated as a positive control.

**Rate of kill assay bioluminescence relative rate of kill (BRRoK) assay.** The assay was performed with *Pf*NF54$^{luc}$ synced trophozoite-stage parasites (20-26h post infection; 2% parasitemia, 2% hematocrit) that were distributed into black-walled clear flat-bottom 96-well test plates at a final volume of 200 μL. Parasites were incubated for 6 h with compounds over a range in concentrations from 30x - 0.1x $EC_{50}$, as determined in a 72h SYBR green assay for *Pf*3D7 ($EC_{50}$ values used were as follows: ART 0.013 μM, DSM265 0.008 μM, **1** 2.1 μM, **2** 0.7 μM, **3** 0.92 μM and **4** 1.9 μM). Bioluminescence was used as a readout for viability at the end of the incubation using the luciferase assay system (Promega) to measure relative light units. After the 6 h incubation, parasites (40 μL) were transferred to 96-well white-walled clear flat-bottom plate. Cells were lysed with passive lysis buffer (10 μL) and luminogenic substrate (50 μL) was added. Bioluminescence was measured on a BioTek Synergy H1 Hybrid plate reader. Assays were performed in technical triplicates on the same plate with two independent biological repeats of each experimental condition. Data were plotted using Graph Pad Prism. DMSO-only wells (parasites + DMSO) were used as a no-kill control to define baseline parasite survival and ART (1.5 μM) was used to define maximal drug-induced killing. Experimental values were normalized to the DMSO and ART controls to generate a relative kill rate dose–response curve, with 100% survival set by the DMSO control and 0% survival defined by the ART total-kill control. Controls included the known fast kill compound ART (defined 100% kill) and the known slow kill compound DSM265, which was not expected to impact cell viability or growth within the 6 h incubation period. [28,34]

***P. falciparum* cytosolic pH assay extended methods.** The assay was performed with *Pf*3D7 synced trophozoite-stage parasites (20-26h post infection) (4% hematocrit, ~5% parasitemia). Infected RBCs (280 mL volume) were distributed into 50 mL conical tubes (30 mL each), pelleted by centrifugation (2500xg for 5 min), resuspended in cold Albumax-free medium (30 mL; RPMI 1640 medium (Millipore Sigma), 23 mM sodium bicarbonate, 92 μM hypoxanthine,12.5 μg/mL gentamicin sulfate, 125 mM NaCl, 25 mM HEPES, 5 mM KCl, 1 mM $MgCl_2$, 20 mM glucose) with the pH adjusted to 7.1 to match the estimated pH of the parasite cytosol and subjected to saponin

(0.1%) lysis of the RBCs. Released parasites were collected by centrifugation at 2500×g for 5 min, supernatant medium containing host cell debris was discarded, and parasite pellets were pooled, resuspended, and washed four times (with 2,500xg, ~5 min centrifugation steps, 50% brake) in an Albumax-free medium (30 mL per wash). Cells were then resuspended at $1\times10^8$ per mL in Albumax-free medium and loaded with the pH-sensitive fluorescent dye ester, BCECF-AM (Thermo Fisher Scientific B1170; dissolved in DMSO at 1 mM) to a final concentration of 1 µM. Cells were then incubated for 30 min at 37°C in 5% $CO_2$. During this time, BCECF-AM permeates into cells and removal of the acetoxymethyl ester by nonspecific esterases traps the non-permeable BCECF dye within the cell. Cells were then washed five times (with 2,500xg, ~5 min centrifugation steps, 50% brake) in "Albumax-free medium" to remove excess extracellular BCECF-AM.

Parasites were then resuspended in ~1 mL (to yield $5.6x10^9$ parasites/mL) "Albumax-free medium" and divided into four centrifuge tubes: 1) $4.2x10^9$ parasites earmarked for 42 experimental wells and placed in a conical tube and 2) 3 microcentrifuge tubes at $1\times10^8$ parasites earmarked for 3 pH controls (pH 6.8, 7.1, 7.8). Parasites were pelleted (conical tube 2,500×g, ~5 min centrifugation steps, 50% brake, microcentrifuge tube 3,000xg ~1 min) and cells in the experimental tube were first washed 2x then resuspended in ~ 9.3 mL pH 7.1 buffer (125 mM NaCl, 25 mM HEPES, 5 mM KCl, 1 mM $MgCl_2$, 20 mM glucose; pH 7.1). These parasites were distributed to 45 wells of a black-walled clear flat-bottom 96-well test plate, yielding $10^8$ parasites per well in a final volume of 198 µL.

Parasites in the three remaining microcentrifuge tubes were washed twice with each individual pH calibration buffer (130 mM KCl, 1 mM $MgCl_2$, 20 mM glucose, 25 mM HEPES; pH 6.8, 7.1, 7.8) and resuspended in 247 µL of each pH buffer respectively. These parasites were distributed to 3 wells of a black-walled clear flat-bottom 96-well test plate, yielding $10^8$ parasites per well in a final volume of 190 µL. Nigericin (Thermo Fisher Scientific N1495; dissolved in

DMSO at 600 μM) was added to the three pH wells (pH 6.8, 7.1, 7.8) (10 μL; 5% final) to generate a standard curve. Under these conditions, the exchange of $H^+$ and $K^+$ ions results in the intracellular pH matching the pH of the external solution. Plates were incubated at 37°C, 5% $CO_2$ for 10 min prior to reading.

For the experimental wells, baseline fluorescence was first established for 10 min, and then CMA (2 μL of a 5 μM stock in 100× DMSO or DMSO vehicle control) was added to 198 μL, resulting in a final concentration of 50 nM contributing 1% DMSO. A new baseline was then established over the next 20 min. Finally, test compounds underwent a standardized serial dilution (40 - 0.002x$EC_{50}$) (8-fold dilution series) in DMSO to 100x of the final assay concentration. Compound (2 μL of 100x DMSO stock or DMSO vehicle control) was added across the experimental wells (final DMSO concentration 2% after addition of both CMA and test compound). Fluorescence measurements as described below were commenced immediately upon addition of experimental compounds and reactions were monitored at 37 °C for 40 min.

Fluorescence was monitored at 37°C on a BioTek Synergy H1 Hybrid plate reader set for 440 nm and 490 nm excitation and 535 nm emission. Two excitation wavelengths were employed; 440 nm was used to monitor pH-independent fluorescence, and then 490 nm monitored the pH-sensitive fluorescence. Emissions were recorded at 535 nm. Variations in the fluorescence ratio (490/440 nm) indicate changes in $pH_{cyt}$. To translate fluorescence ratio readings into $pH_{cyt}$ values, fluorescence ratio readings for the pH wells (pH 6.8, 7.1, 7.8) taken over the total time of 70 min were averaged and plotted as the x-axis against the pH of the solution 6.8, 7.1, 7.8 y-axis to fit a straight-line relationship. A regression line was fitted to the data (fluorescence ratio = m × $pH_{cyt}$ + c), where m and c represent the slope and y-intercept, respectively and the calculated values for m and c were utilized for the conversion of experimental fluorescence ratio readings into $pH_{cyt}$ values.

***P. falciparum* pH fingerprint assay (detailed methods).** Studies were performed as previously described.[41] Briefly, trophozoite-stage *P. falciparum* parasites (3D7 strain) were isolated from their host erythrocytes via brief exposure to saponin, loaded with the pH-sensitive fluorescent dye BCECF, then incubated for 20 min in a glucose-free saline (135 mM NaCl, 5 mM KCl, 1 mM $MgCl_2$, 25 mM HEPES; pH 7.10) to deplete ATP. Parasites were then added to three different solutions (to which a test compound, control compound, or solvent alone was added): (**1**) glucose-containing saline solution (125 mM NaCl, 5 mM KCl, 1 mM $MgCl_2$, 20 mM glucose, 25 mM HEPES; pH 7.1), (**2**) glucose-containing saline solution with the V-type $H^+$-ATPase inhibitor concanamycin A (final concentration 100 nM; added from a 100 μM DMSO stock introducing 0.1% v/v DMSO), and (**3**) a saline solution lacking glucose and $Cl^-$ (135 mM $Na^+$-gluconate, 5 mM $K^+$-gluconate, 1 mM $MgSO_4$, 25 mM HEPES; pH 7.10), creating the 'low Cl- condition' (final external [$Cl^-$] in the assay = 14.2 mM). Fluorescence was monitored at 37˚C for 40 min using excitation wavelengths of 440 nm and 495 nm and an emission wavelength of 520 nm. Fluorescence Ratio values (495 nm/440 nm) were converted to $pH_{cyt}$ as previously described.[44] Control compounds were the *Pf*ATP4 inhibitor KAE609 (50 nM), the protonophore CCCP (100 nM), the *Pf*FNT inhibitor MMV007839 (2 μM), the non-specific $Cl^-$ transport inhibitor DIDS (sodium (E)-6,6'-(ethene-1,2-diyl)bis(3-isothiocyanatobenzenesulfonate) (100 μM), the *Pf*HT inhibitor MMV009085 (5 μM), the V-type $H^+$-ATPase inhibitor concanamycin A (100 nM) and DMSO (0.1% v/v; solvent control). Compounds **2** and **3** were tested at a concentration of 5 μM. Compounds were diluted to their final concentrations in the assay from DMSO stocks, introducing 0.1% v/v DMSO.

***In vitro* ADME (extended methods).** For solubility measurements, compounds were dissolved in DMSO and spiked into phosphate buffer (pH 6.5) or 0.01 M HCl (approx. pH 2.0). The final DMSO concentration was 1%. After 30 minutes, solubility was determined by nephelometry analysis.[17] For human liver microsomes, 1 µM compound was incubated with human liver microsomes (Xenotech, lot# 1410230, 0.4 mg/mL protein) at 37 ºC. An NADPH-regenerating system was added to initiate metabolism; control reactions did not contain NADPH.[17] At various time points over the course of an hour, samples were collected and quenched using acetonitrile, with diazepam included as an internal standard. In vitro intrinsic clearance ($CL_{int}$) was calculated from the apparent first-order degradation rate constant.

**Supplemental Tables**

**Table S1.** Commercially sourced analogs of SW491 (**2)** and SW968 (**3**) were evaluated for activity against *P. falciparum* parasites

A. **2**-analogs

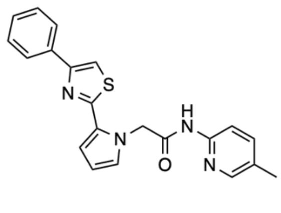

SW463 (**5**)

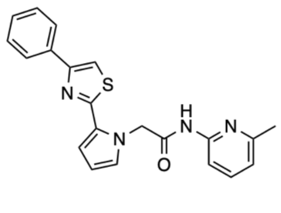

SW316 (**6**)

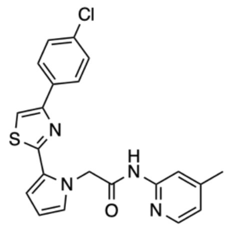

SW317 (**7**)

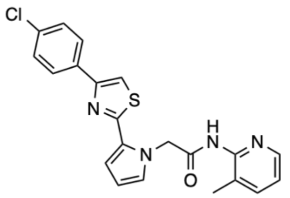

SW318 (**8**)

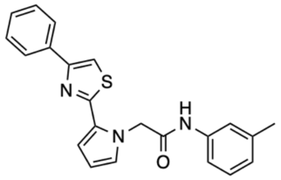

SW319 (**9**)

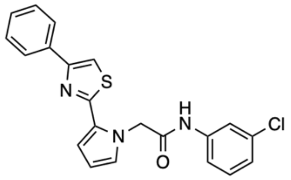

SW320 (**10**)

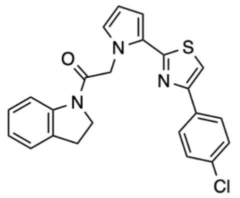

SW966 (**11**)

| Parasite | **5** | **6** | **7** | **8** | **9** | **10** | **11** |
|---|---|---|---|---|---|---|---|
| *Pf*Dd2 $EC_{50}$ (µM) | 9.6 ± 0.44 (2) | 12 ± 1.1 (2) | 14 ± 0.58 (2) | >30 (1) | 1.7 ± 0.038 (2) | 0.98± 0.082 (2) | >30 (1) |
| *Pf*3D7 $EC_{50}$ (µM) | 10 ± 0.030 (2) | 15 ± 0.87 (2) | 15 ± 2.4 (2) | >30 (1) | 2.8 ± 0.17 (2) | 1.6 ± 0.33 (2) | >30 (1) |

**2**-analogs were at least 95% pure as assessed by LC/MS. *P. falciparum* $EC_{50}$ values represent the mean ± standard error from two independent biological replicates, with the number of independent replicates in parenthesis, with exception for **11** and **8**, where only one study (3 technical replicates) was performed.

B. **3**-analogs

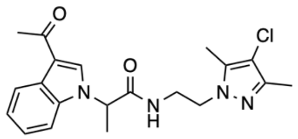

SW181 (**12**)

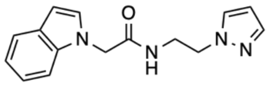

SW314 (**13**)

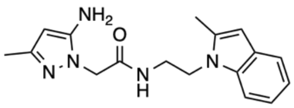

SW315 (**14**)

| Parasite | **12** | **13** | **14** |
|---|---|---|---|
| *Pf*Dd2 $EC_{50}$ (μM) | >30 (1) | >30 (1) | >30 (1) |
| *Pf*3D7 $EC_{50}$ (μM) | >30 (1) | >30 (1) | >30 (1) |

**3**-analogs were at least 95% pure, as assessed by LC/MS. *P. falciparum* $EC_{50}$ values showed no activity at 30 μM. As a result, only a single biological experiment was conducted for each compound, with three technical replicates.

**Table S2.** Summary of *Pf*3D7 cellular and pH $EC_{50}$ data for DSM265 and KAE609

| | DSM265 | KAE609 |
|---|---|---|
| Activity of all compounds on *Pf* and human cells (nM) | | |
| *Pf*3D7 $EC_{50}$ | 6.6 ± 0.36 (3) | 1.0 ± 0.038 (3) |
| *Pf*3D7 pH $EC_{50}$ | >40-fold $EC_{50}$ (2) | 3.0 ± 2.1 (2) |

*P. falciparum* $EC_{50}$ values (72 h Sybr green assay) represent the mean ± standard error with the number of independent replicates in parentheses. *P. falciparum* pH $IC_{50}$ values represent the mean ± std deviation from two independent biological replicates, see Figure 2D and Figure S2D.

**Table S3.** Resistance selection recrudescence day and bulk culture $EC_{50}$ data for SW491 (**2**) and SW968 (**3**)

| Time to Recrudescence | | | | | |
|---|---|---|---|---|---|
| | Pulse 1 10×$EC_{50}$ | Pulse 2 10×$EC_{50}$ | Pulse 3 20×$EC_{50}$ | Pulse 4 20×$EC_{50}$ | Pulse 5 20×$EC_{50}$ |
| **2**-selected, Flasks #1-4 | 8 days | 6 days | 7 days | 3 days | 48 hours after pulse |
| **3**-selected, Flasks #1-4 | 5 days | 4 days | 4 days | 6 days for Flask #1; 2 days for Flasks 2 - 4 | |
| SW491 (**2**) and SW968 (**3**) $EC_{50}$ (μM) | | | | | |
| | Pulse 1 | Pulse 2 | Pulse 3 | Pulse 4 | Pulse 5 |
| **2**-selected Flask 1 | nd | nd | 0.57 | 8.4 | nd |
| **2**-selected Flask 2 | nd | nd | 0.58 | >10 | nd |
| **2**-selected Flask 3 | nd | nd | 1.1 | >10 | nd |
| **2**-selected Flask 4 | nd | nd | 0.81 | 6.6 | nd |
| **3**-selected Flask 1 | nd | 0.72 | 1.2 | nd | n/a |
| **3**-selected Flask 2 | nd | 0.84 | >10 | nd | n/a |
| **3**-selected Flask 3 | nd | 1.0 | >10 | nd | n/a |
| **3**-selected Flask 4 | nd | 1.1 | >10 | nd | n/a |
| **2** Dd2 Parent (clone B8) | nd | nd | 0.78 | 0.78 | nd |
| **3** Dd2 Parent (clone G9) | nd | 0.99 | 0.99 | nd | n/a |

Resistance selections for **2** and **3** using *Pf*Dd2 parasites: Upper panel, Time to recrudescence after each pulse; Lower panel, $EC_{50}$ values of bulk cultures after indicated rounds of selection. nd – not determined n/a – not applicable

**Table S4.** Protein coding mutations observed in **2** and **3** resistant parasites

A. Mutations observed in *Pf*ATP4

| **Strain** | ***Pf*ATP4 (PfDd2_120016700)** |
|---|---|
| **2**-selected | |
| Dd2 parent (clone B8) | P412 (CCT), F917 (TTC) |
| **2**-resistant Flask 1 Clone B6 | P412 (CCT), L917 (TT**A**) |
| **2**-resistant Flask 2 Clone F9 | L412 (C**T**T), F917 (TTC) |
| **2**-resistant Flask 3 Clone F5 | L412 (C**T**T), F917 (TTC) |
| **2**-resistant Flask 4 Clone E10 | L412 (C**T**T), F917 (TTC) |
| **3**-selected | |
| Dd2 parent (clone G9) | F917 (TTC) |
| **3**-resistant Flask 2 Clone D10 | L917 (TT**A**) |
| **3**-resistant Flask 3 Clone D8 | L917 (TT**A**) |
| **3**-resistant Flask 4 Clone B3 | L917 (TT**A**) |

WGS detected mutations for *Pf*ATP4 (PfDd2_120016700) (SNPs, red) and these changes were confirmed by Sanger sequencing in all parental and clonal lines from the Dd2 screen. Figures S6-S7. WGS data are available under (SRA) database (SRA BioProject ID PRJNA1230203).

B. Polymorphisms observed in additional genes found in some but not all clones

| Strain | ***Pf*S8E (PfDd2_070012100)** | ***Pf* Conserved protein (PfDd2_130068800)** |
|---|---|---|
| Dd2 parent (clone B8) | L62 | Y567 |
| **2**-selections | | |
| **2**-resistant Flask 1 Clone B6 | **R**62 | Y567 |
| **2**-resistant Flask 2 Clone F9 | L62 | Y567 |
| **2**-resistant Flask 3 Clone F5 | L62 | Y567 |
| **2**-resistant Flask 4 Clone E10 | L62 | Y567 |
| **3**-selections | | |
| Dd2 parent (clone G9) | L62 | Y567 |
| **3**-resistant Flask 2 Clone D10 | L62 | **S**567 |
| **3**-resistant Flask 3 Clone D8 | L62 | **S**567 |
| **3**-resistant Flask 4 Clone B3 | L62 | **S**567 |

WGS detected SNPs (SNPs, red) for *Pf*S8E (PfDd2_070012100) and *Pf* Conserved protein (PfDd2_130068800), and *Pf*MDR1 (PfDd2_050027900) compared to the parental line.

**Table S5:** Cross resistance data for additional clonal lines of **2** and **3** resistant parasites.

| Parasite line | Resistance | SW412 (**1**) (μM) | SW491 (**2**) (μM) | SW968 (**3**) (μM) | SW080 (**4**) (μM) | KAE609 (nM) |
|---|---|---|---|---|---|---|
| Dd2 | CQ, CYC, PYR | 2.4[a] | 0.83 [a] | 1.1 [a] | 1.6 [a] | 1.2 [b] |
| **2**-resistant Flask 1 Clone B6 (Clone B8) | $PfATP4^{F917L}$ | >10 (3) | 9.6 ± 2.0 (3) | >10 (3) | >12 (2) | 12 ± 0.98 (3) |
| **2**-resistant Flask 2 Clone F9 (Clone B8) | $PfATP4^{P412L}$ | 7.5 ± 2.4 (3) | >10 (3) | >10 (3) | 4.0 ± 0.090 (2) | 14 ± 1.4 (3) |
| **2**-resistant Flask 3 Clone F5 (Clone B8) | $PfATP4^{P412L}$ | 7.6 ± 2.0 (3) | >10 (3) | >10 (3) | 3.2 ± 0.13 (2) | 12 ± 1.4 (3) |
| **2**-resistant Flask 4 Clone E10 (Clone B8) | $PfATP4^{P412L}$ | 9.4 ± 2.5 (3) | >10 (3) | >10 (3) | 4.5 ± 0.14 (2) | 14 ± 1.1 (3) |
| **3**-resistant Flask 2 Clone D10 (Clone G9) | $PfATP4^{F917L}$ | >10 (3) | >10 (3) | >10 (3) | 11 ± 1.7 (2) | 12 ± 0.34 (3) |
| **3**-resistant Flask 3 Clone D8 (Clone G9) | $PfATP4^{F917L}$ | >10 (3) | >10 (3) | >10 (3) | 10 ± 0.34 (2) | 12 ± 0.97 (3) |
| **3**-resistant Flask 4 Clone B3 (Clone G9) | $PfATP4^{F917L}$ | >10 (3) | >10 (3) | >10 (3) | >12 (2) | 14 ± 0.90 (3) |

Data were reproduced from [a] Table 1 or [b] Table 2. Data represent the mean ± standard error from independent experiments with the number of replicates shown in parenthesis. Each independent study is derived from triplicate technical replicates. Top concentrations in the dose-response titrations were 10μM for **1**-**3**, 12μM for **4** and 0.05 – 0.25 μM for KAE609. (Clone B8) and (Clone G9) refer to Dd2 parent clones.

**Table S6:** Primers for PCR amplification and Sanger sequencing of *pfatp4, pfrs8e, pfcuf, and pfmdr1.* (Phillips Lab)

| Gene | Positions of interest | Primers |
|---|---|---|
| *pfatp4* (PfDd2_120016700) | 412 (P412) | F: 5'-CTGAACAAGTAAAAATAAATAGAGACA-3'<br>R: 5'-TTCCTTCAGTTAATGTACCGG-3' |
| *pfatp4* (PfDd2_120016700) | 917 (F917) | F: 5'-TGGAGTTAATGATGCACCTGC-3'<br>R: 5'-CATCATTTGGTGGTTCTCTTG-3' |
| Ribosomal protein S8e, putative *pfrs8e* (*Pf*Dd2_070012100) | 63 (K63) | F: 5'- GGATATCGCTTTGATCATTTCGAA -3'<br>R: 5'- TGGGTAGTTGCCATTTTCCAC -3' |
| conserved Plasmodium protein, unknown function *pfcpuf* (*Pf*Dd2_130068800) | 567 (Y567) | F: 5'- GAAGGACATTCTTTTTTTGGCTAG-3'<br>R: 5'- CAAGTCAAAATGTTCCCCTTC -3' |
| multidrug resistance protein 1 *pfmdr1* (PfDd2_050027900) | 86 (F86Y) | F: 5'-GAGTACCGCTGAATTATTTAGAA-3'<br>R: 5'-TTATTATCATGAAATTGTCCATCTTG-3' |

**Table S7:** Primers used for PCR amplification and sequencing of *pfatp4* (PF3D7_1211900) (Fidock Lab).

| Primer Name | Sequence (5'→ 3') | PCR Function |
|---|---|---|
| P6536 | ATGAGTTCTCAAAATAATAATAAACAGGGTGGAC | Outer Flank PCR, Sequencing |
| P6537 | TTAATTCTTAATAGTCATATATTTTCTTCTATATATAACCTTTGG | Outer Flank PCR, Sequencing |
| P8180 | ATTCATTAAAAAATGATGAATTAAATAAAAATACAACGATG | Nested, Sequencing |
| P8181 | TTGCCACCATAACAAACATGTTGTATTCAAAATA | Nested, Sequencing |
| P6538 | TATTCAAGAGCACAACCGGAAG | Sequencing |
| P6539 | GCATTATGTGTCTTGTTATCATTGGC | Sequencing |
| P6540 | GGGTACTTCTATCAAGTAATCTATCAGGTGC | Sequencing |
| P6541 | GCTGTATCTTCCATTCCAGAAGG | Sequencing |
| P6560 | ACCTTGAATGCTTGCTTAGCAACC | Sequencing |
| P6561 | CGAGAATGTATATTTAGATAAACCTGG | Sequencing |
| P6562 | TCGAGACGGTATAACTACCTTCTGACC | Sequencing |
| P6563 | GCATCCTAAAGTTTCAACAGCTGGTAG | Sequencing |
| P6564 | TCTGTTCCATTAATACCCATAGCAACAC | Sequencing |
| P8175 | TAATTCCAATAATGTAGAAGAC | Sequencing |
| P8176 | AATGCCATTCAAGTTATAAAAAC | Sequencing |

**Table S8:** Characteristics of the Dd2 *Pf*ATP4 mutant lines used for cross-resistance profiling (Fidock Lab). This table shows the parasite line name, *Pf*ATP4 genotype, and $EC_{50}$ values (μM) and corresponding $EC_{50}$ fold-changes compared to the Dd2-B2 parent (drug-sensitive/wild-type *Pf*ATP4) when tested for susceptibility to the *Pf*ATP4 inhibitors, KAE609 (spiroindolone), and the dihydroisoquinolone analogs, **15** and SJ733.

| Parasite Line ID (clone ID) | *Pf*ATP4 Genotype | Compound | | | | | |
|---|---|---|---|---|---|---|---|
| | | SJ733 | | MMV609 (**15**) | | KAE609 | |
| | | [a]$EC_{50}$ (μM) | [b]Fold change | [a]$EC_{50}$ (μM) | [b]Fold change | [a]$EC_{50}$ (μM) | [b]Fold change |
| Dd2-B2 | wildtype | 0.13± 0.012 (2) | 1 | 0.0066± 0.00030 (4) | 1 | 0.0021± 0.00030 (2) | 1 |
| [c]$Dd2^{G358S}$ (Dd2-SJ16-D2) | $ATP4^{G358S}$ | [e]31 (1) | 240 | ND | ND | 1.5± 0.11 (2) | 710 |
| [d]$Dd2^{L350H}$ (MMV609 10×$IC_{50}$_fl2_E3) | $ATP4^{L350H}$ | ND | ND | 0.20 ± 0.071 (2) | 30 | [f]0.0048 | [f]3.7 |
| [d]$Dd2^{P412L}$ (MMV609 10×$IC_{50}$_fl2_C11) | $ATP4^{P412L}$ | ND | ND | 12 ± 4.5 (2) | 1800 | [f]0.011 | [f]8.5 |

[a] $EC_{50}$ values are presented as means ± standard error of the mean. The number of independent experiments are shown in parenthesis and each was derived from technical duplicates.
[b] Fold-changes in the mean $EC_{50}$ values of the PfATP4 mutant lines were determined for each *Pf*ATP4 mutant line compared to the Dd2-B2 parent.
[c] $Dd2^{G358S}$ is a SJ733-selected clone (DD2-SJ16-D2) that expresses the *Pf*ATP4 G358S mutation and was reported in a previous study to cause a 5-fold $EC_{50}$ increase against SJ733.[26] Our results identify a much higher level of resistance, and they match well with data reported in a second study that found that this mutation led to very high levels of resistance for both SJ733 and KAE609.[21]
[d] $Dd2^{L350H}$ and $Dd2^{P412L}$ are **15**-selected clones expressing the *Pf*ATP4 L350H and P412L mutations, respectively.
[e] The $EC_{50}$ value of $Dd2^{G358S}$ when tested against SJ733 was estimated from one biological replicate, performed in duplicate. This fold shift was substantially higher than the earlier report of a 5-fold shift.
[f] Data in this table represent an independent data set from Table 3, except for $Dd2^{L350H}$ and $Dd2^{P412L}$ mutant lines versus KAE609 where the data are reproduced from Table 3. The fold-changes in the mean KAE609 $EC_{50}$ values shown for the $Dd2^{L350H}$ and $Dd2^{P412L}$ mutant lines are compared to an $EC_{50}$ value of 0.0013 μM for the Dd2-B2 parent when tested against KAE609 in the latter dataset (Table 3).
ND, not determined.

**Supplemental Figures**

**Figure S1.**

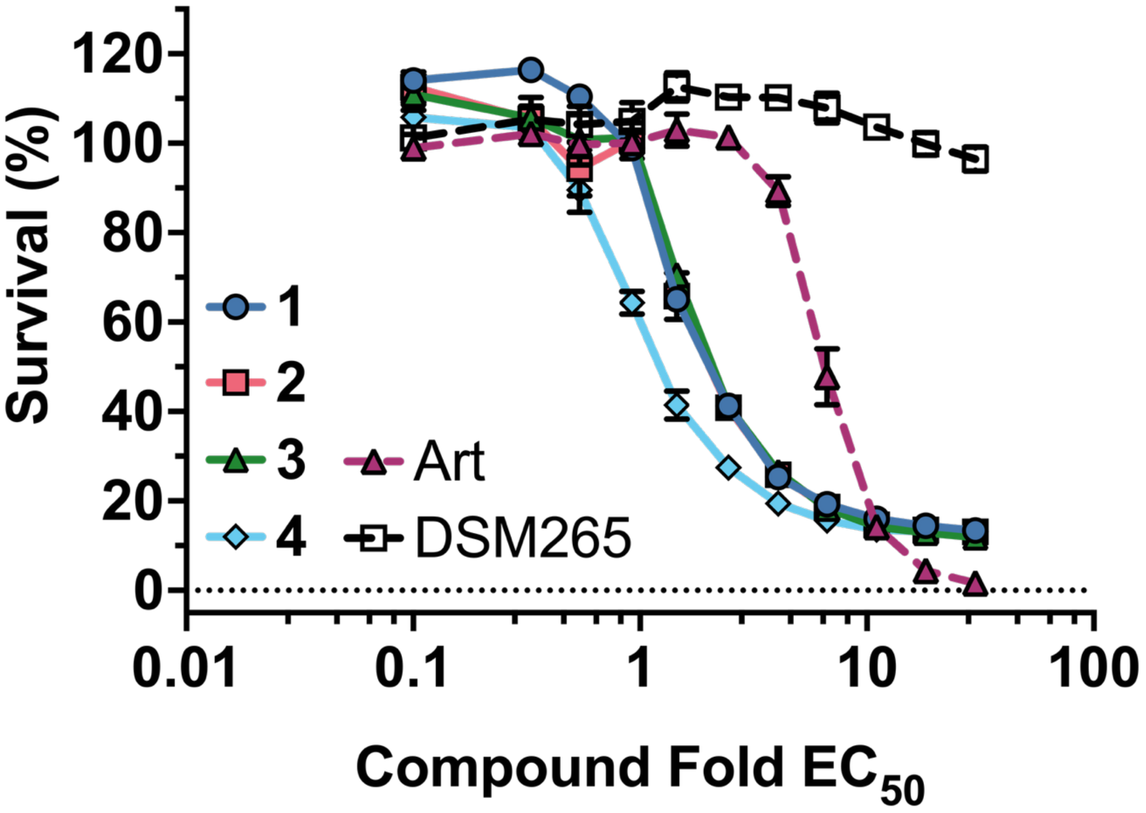


**Fig. S1.** Additional replicate of α-azacyclic acetamides kill rate data in support of Figure 1.

Kill rate was assessed using the BRRoK assay. These data represent a second independent replicate for the data reported in Figure 1C. Data are the mean ± std dev for 3 technical replicates. Refer to Figure 1C caption for additional information on experimental design.

**Figure S2.**

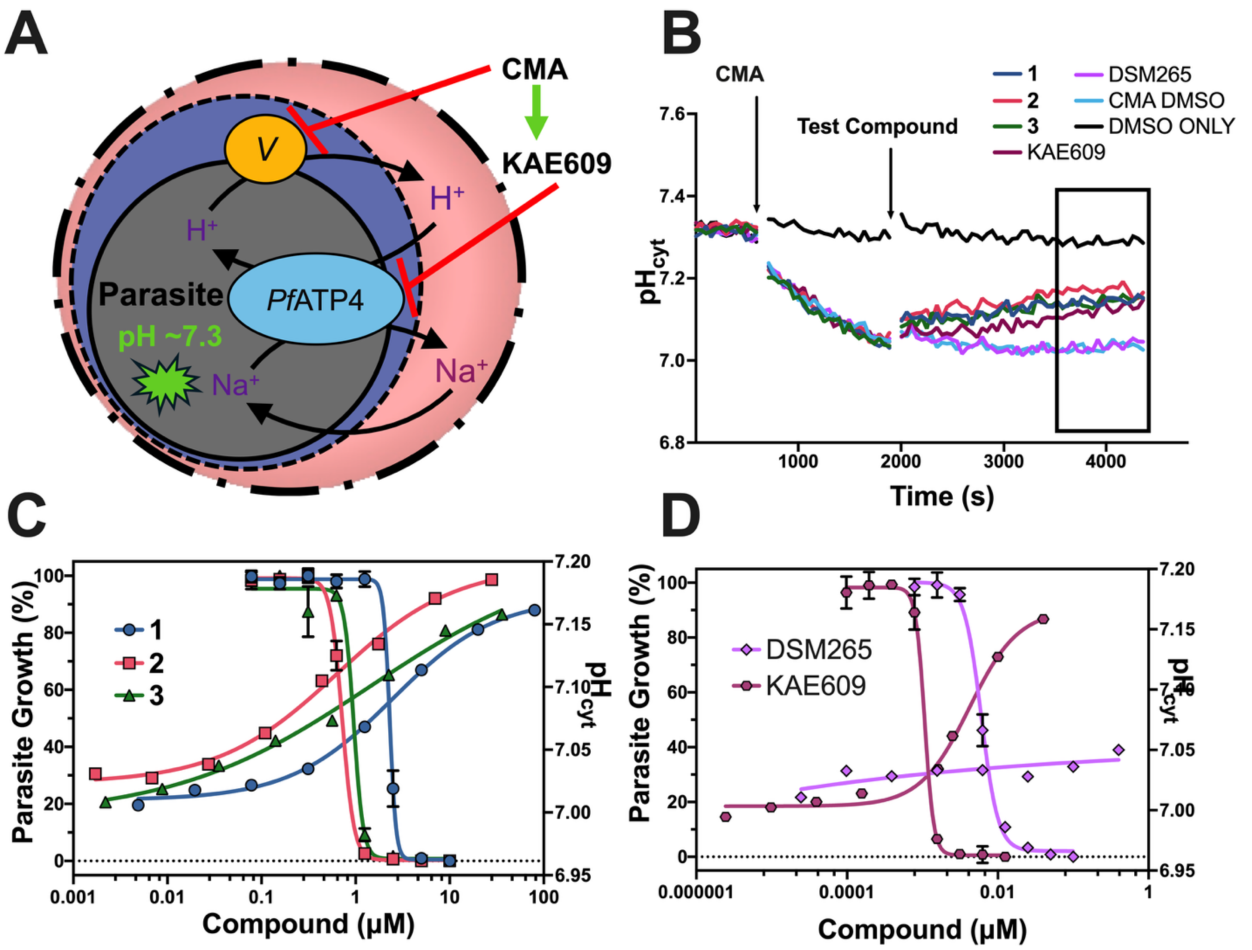


**Figure S2.** The effects of **1**-**3** on intracellular pH measured in a second independent experiment in support of Figure 2.

(A) Schematic representation of *Pf*ATP4 (reproduced from Figure 2). (B) a representative $pH_{cyt}$ versus time curve showing compound-induced changes at a compound concentration of $10xEC_{50}$. (C) The average $pH_{cyt}$ value obtained from the final 10-minute period of the assay is plotted versus compound concentration (right hand axis) and a representative dose response for parasite growth reproduced from Figure 2 is plotted on the left axis. (D) Similar graphical representation and $EC_{50}$ overlay as in (C) for controls DSM265 and KAE609. These pH studies represent a second independent replicate to the studies described in Figure 2. Refer to Figure 2 caption for additional information on experimental design.

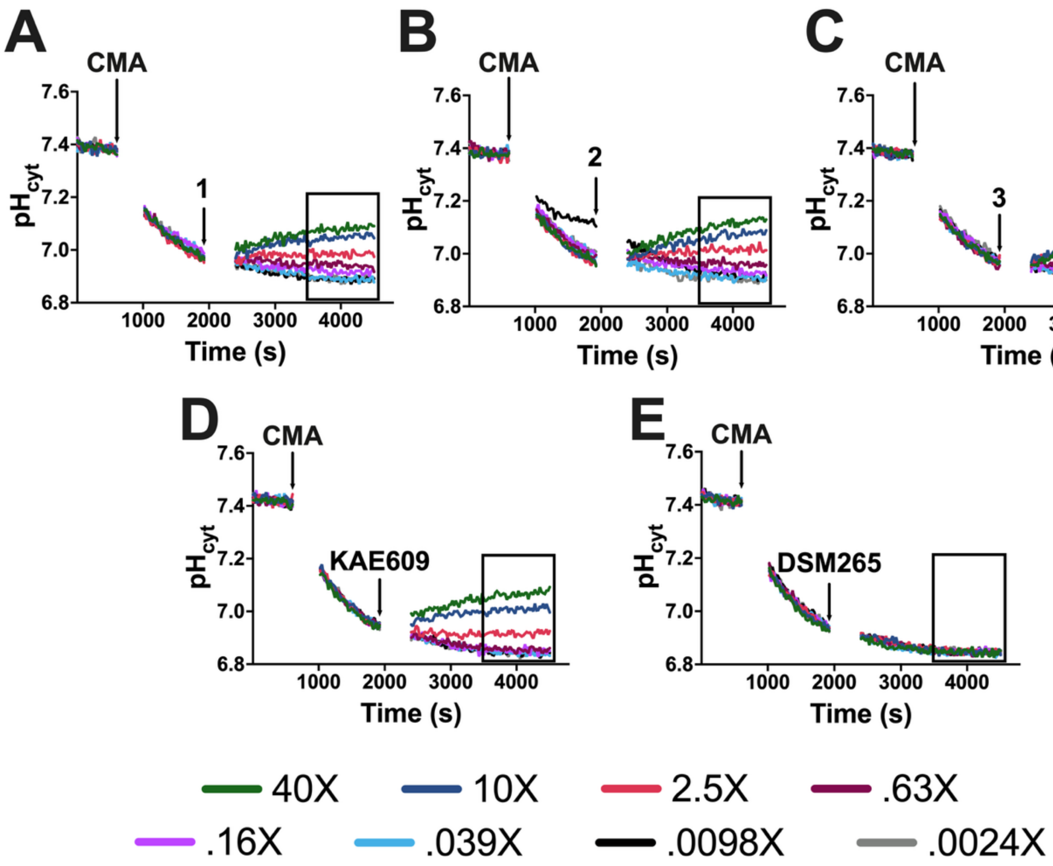


**Figure S3.** Full pH versus time profiles showing the effects of a range of concentrations of α-azacyclic acetamides **1-3** (A-C) and control compounds (E, F) on intracellular pH.

(A-E) Representative data from a single experimental trial are presented for a range of compound concentrations (40–0.002x$EC_{50}$) using an 8-point dilution series (strain 3D7), as outlined in Figure 2B. The average pH values were calculated over a 10-minute interval corresponding to the period of maximal acidification observed for the controls (DSM265 and Concanamycin A (CMA); see highlighted boxes). The 10x$EC_{50}$ curves from these plots are reproduced in Figure 2B.

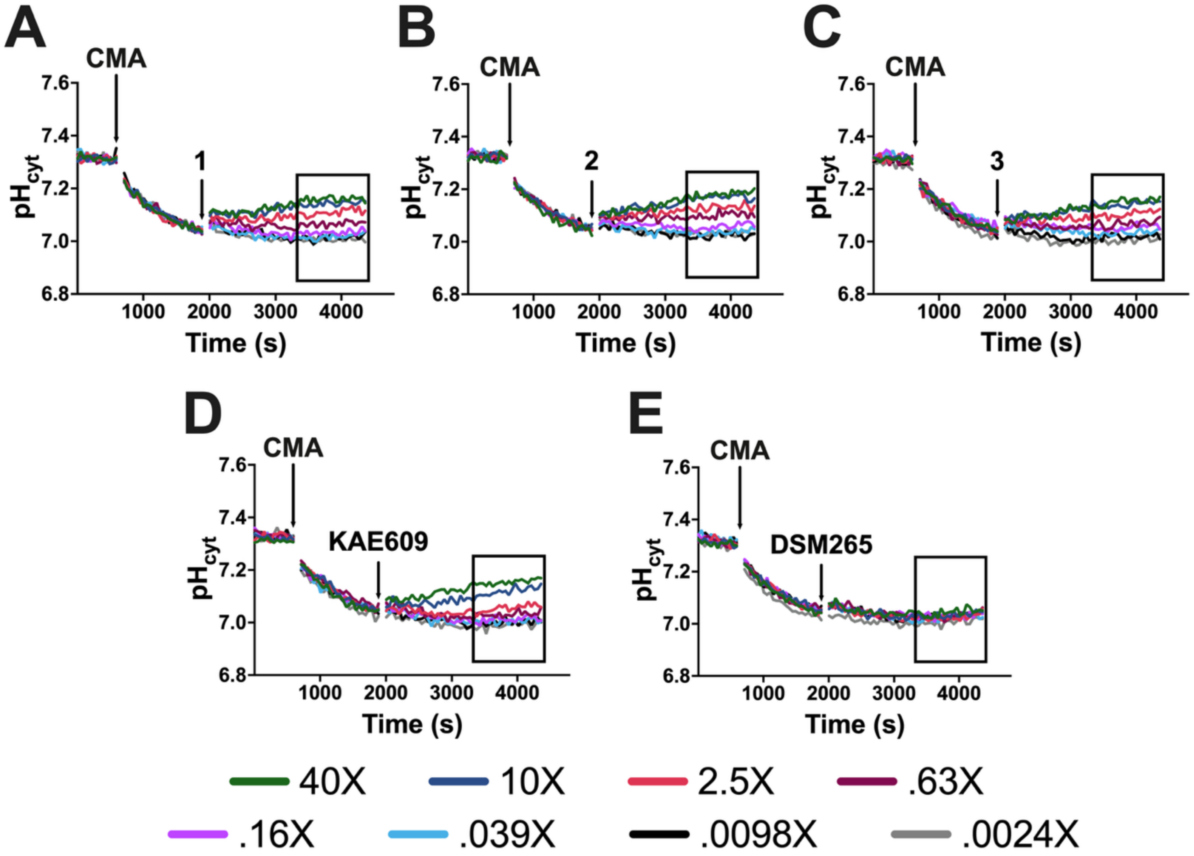


**Figure S4.** Full pH versus time profiles showing the effects of a range of concentrations of $\alpha$-azacyclic acetamides **1-3** (A-C) and control compounds (E, F) on intracellular pH. This is a second experimental replicate to data shown in Figure S3 and this figure supports data shown in Figures S2C and S2D.

(A-E) Representative data from a single experimental trial are presented across a range of compound concentrations (40–0.002x$EC_{50}$) using an 8-point dilution series (strain 3D7), as outlined in Figure S2B. The average pH values were calculated over a 10-minute interval corresponding to the period of maximal acidification observed for the controls (DSM265 and Concanamycin A (CMA); see highlighted boxes). The 10x$EC_{50}$ curves from these plots are reproduced in Figure S2B.

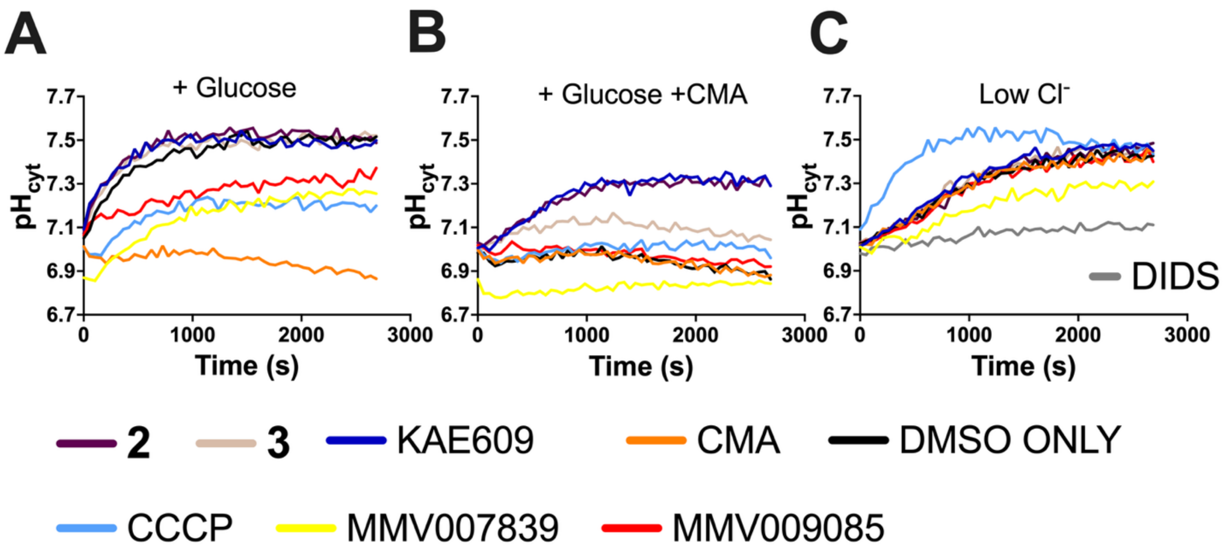


**Figure S5.** *Pf*ATP4 pH fingerprint assay.

(A-C) *Pf*3D7 trophozoites isolated from their host erythrocytes by brief exposure to saponin were loaded with the pH-sensitive fluorescent dye BCECF and placed in a glucose-free saline solution for 20 minutes to deplete ATP. $pH_{cyt}$ was then monitored over time in parasites that were added to: (**A**) a saline solution containing glucose, (**B**) a saline solution containing glucose and the V-type $H^+$-ATPase inhibitor concanamycin A (CMA; 100 nM), and (**C**) a saline solution containing no glucose in which $Cl^-$ has been replaced with gluconate. $\alpha$-azacyclic acetamides **2** and **3** were both tested at 5 $\mu$M. The controls are the *Pf*ATP4 inhibitor KAE609 (50 nM), the protonophore CCCP (100 nM), the *Pf*FNT inhibitor MMV007839 (2 $\mu$M), the non-specific $Cl^-$ transport inhibitor DIDS (100 $\mu$M), the *Pf*HT hexose transporter inhibitor MMV009085 (5 $\mu$M), the V-type $H^+$ ATPase inhibitor CMA (100 nM), and DMSO (0.1% v/v; solvent control). The data are from a single experiment, representative of two similar experiments in which **2** and **3** were tested.

**Figure S6.** Sanger sequencing to verify **2** (SW491)-selected resistance mutations in *pfatp4* (PfDd2_120016700)

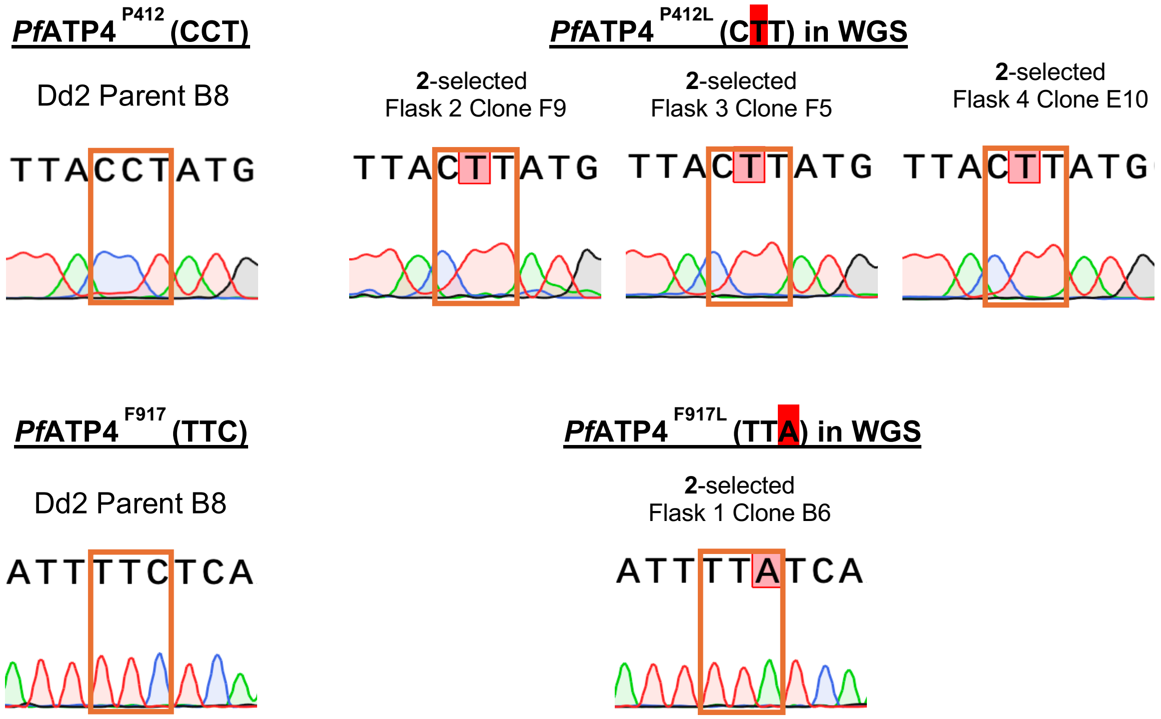


**Figure S7.** Sanger sequencing to verify **3** (SW968)-selected resistance mutations in *pfatp4* (PfDd2_120016700)

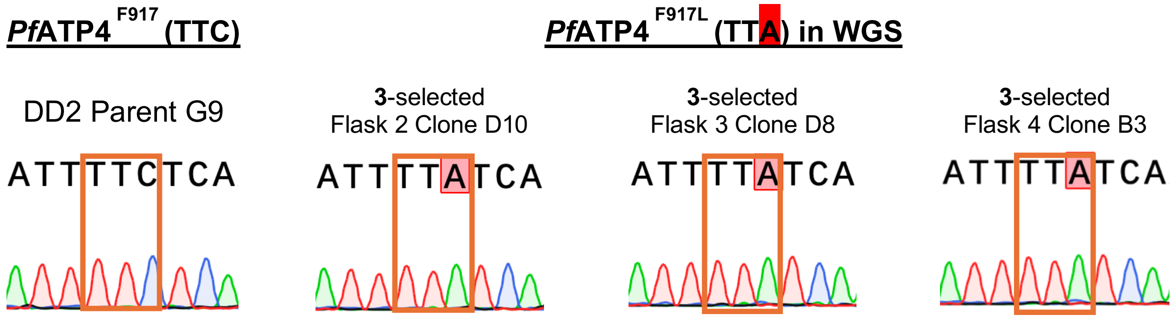

**Figure S8.** Long read sequencing to verify **2**-selected mutations in Ribosomal protein S8e, putative *pfrps8e* (*Pf*Dd2_070012100)

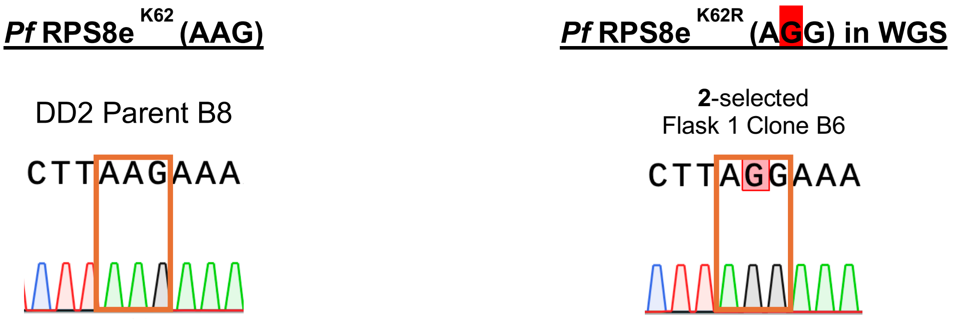


**Figure S9.** Long-read sequencing to verify **3**-selected mutations in conserved Plasmodium protein, unknown function *pfcpu* (*Pf*Dd2_130068800)

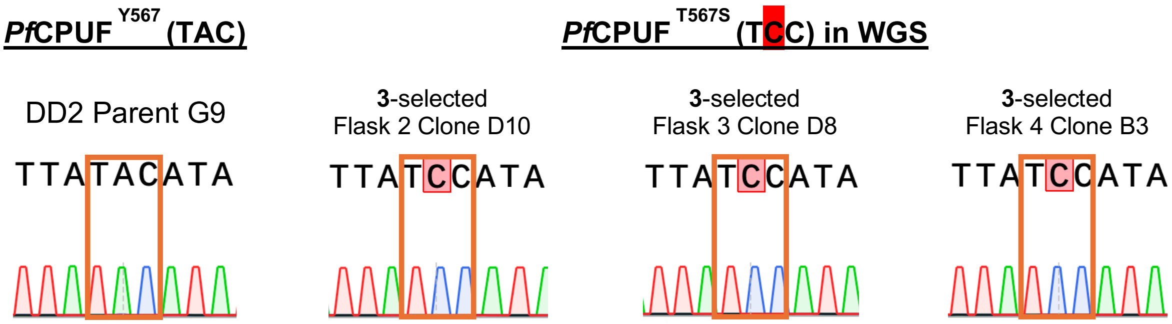


**Figure S10.** Long-read sequencing of multidrug resistance protein 1 *pfmdr1* (*Pf*Dd2_050027900) revealed a mixture of F86 alleles (blue box) in the parental Dd2 lines and **3**-selected clonal mutant lines.

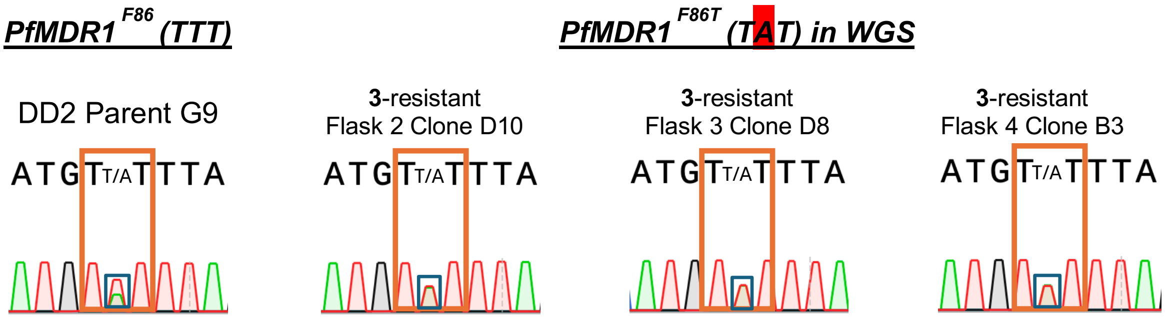
